# The effect of environmental enrichment on immune cell DNA methylation profiles depends on the parity of sows

**DOI:** 10.1101/2025.07.01.662510

**Authors:** Mariana Mescouto Lopes, Gabriel Costa Monteiro Moreira, Aurélie Chaulot-Talmon, Anne Frambourg, Valentin Costes, Julie Demars, Elodie Merlot, Hélène Jammes

**Affiliations:** PEGASE, INRAE, Institut Agro, 35590 Saint-Gilles, France; BREED, Université Paris-Saclay, UVSQ, INRAE, 78352 Jouy-en-Josas, France; ELIANCE, 149 rue de Bercy, 75595 Paris cedex 12, France; GenPhySE, INRAE, Université de Toulouse, 31326 Castanet Tolosan, France

**Keywords:** Epigenetics, Biomarkers, Pig, Reproductive life, Positive welfare, Immunity

## Abstract

The aim of this study was to identify epigenetic markers that reflected positive affective states in multiparous pregnant sows. The animals were housed during gestation in either a conventional (C) environment (2.4 m² per sow), featuring a concrete slatted floor and minimal enrichment, or in an enriched (E) environment (3.5 m² per sow) with deep straw bedding. Peripheral blood mononuclear cells (PBMCs) were isolated at day 98 of gestation (G98) and day 12 of lactation (L12) for genomic DNA extraction and reduced representation bisulfite sequencing (RRBS). Significant effects of individual class of parity (low-parity (LP) versus high-parity (HP)) were observed with the identification of 1,358 and 680 differentially methylated cytosines (DMCs) at G98 and L12, respectively. Interestingly, some of them are organized into differentially methylated regions (DMRs). Some DMRs colocalized with or near a gene and displayed a continuous methylation distribution (44 at G98 and 15 at L12 including 4 in common): 5 targeted genes are related to epigenetic regulation (*DNMT3A*, *KDM8*, *HDAC4*, *SIRT2* and *U2*) and 12 to immune function (*CD2*, *CD5*, *CAMK4*, *SECTM1*, *URODL1*, *CMIP*, *SEC14L1, SKI*, *TNFRSF1B*, *CCND3*, *SGK1*, *PACS1*). These results suggested a true epigenetic impact of parity class on individual immunity. Considering the two parity classes separately, a minor effect of environmental enrichment was observed (at G98: 60 and 42 DMCs; at L12: 35 and 81 DMCs, in the LP and HP groups, respectively). Remarkably, some DMC-associated genes had previously been linked to affective states in humans. In conclusion, unexpected DNA methylation changes associated with parity class were identified, suggesting a genome adaptation during reproductive life and modifying the response to housing conditions. Furthermore, specific CpG sites emerged as potential biomarkers of positive affective states in pigs.

**Highlights:**

- This study revealed a substantial impact of the genetic background of sows on DNA methylation patterns, emphasizing the need to account for genetic factors when analysing epigenetic data.
- Parity, which was confounded with the age of the sows in this study, significantly influenced DNA methylation, even among multiparous sows, highlighting the importance of this factor in modulating epigenetic mechanisms in immune cells.
- We observed minor differences in the effects of environmental enrichment on DNA methylation profiles, which differed depending on the parity group.
- We suggest a short list of possible biomarkers with biological meaning responding to environmental enrichment.

## Introduction

Affective states, or the subjective positive or negative feelings experienced by animals, play a central role in their welfare^1–3^. Given the inability of non-human animals to communicate their feelings verbally, scientists are reliant on indirect behavioural and physiological tests to assess their affective states^4.^ While tools are now available to identify responses to acute emotions, it is more challenging to assess long-term, and usually less intense, emotions (i.e., moods). Animal welfare science would therefore benefit from the development of reliable, feasible, and practical physiological markers that reflect the affective states of animals.

In mainstream pig production systems, one of the primary sources of animal stress is the barren housing environment in which they are raised. The lack of adequate stimuli limits the expression of natural behaviours and cognitive-sensory capacities, leading to signs of chronic stress and poor welfare. Environmental enrichment has been shown to positively impact pig behaviour^5,6^, cognitive function^7,8^, and physiological variables such as cortisol levels^9,10^. Enriched housing conditions have also been found to modulate the blood immune response and reduce disease susceptibility in pigs^11,12^. Blood immune cells may therefore offer valuable insights into the influence of living environments on the affective state of animals.

Epigenetic mechanisms involve stable and heritable modifications during cell division, which regulate gene expression without altering nucleotide sequences^13^. These mechanisms play a crucial role in development and cell differentiation and enable the integration of environmental factors to shape cellular function. For instance, DNA methylation, which encompasses the addition of a methyl (CH3) group to a cytosine residue within a CG dinucleotide, is essential to establishing and preserving cellular identity. During haematopoiesis, from stem cells to highly differentiated immune cells, specific DNA methylation profiles are established and support the unique functions of each immune sub-type cells^14,15^. Despite this specificity, DNA methylation in blood immune cells can be influenced by various internal and external factors throughout an individual’s life, from early embryonic development to death. In humans and rodents, changes of DNA methylation patterns in blood immune cells have been associated with age^16^, physiological stages^17^, toxic exposure^18^ and prenatal stress^19^.

In pigs, specific DNA methylation profiles have been described for different immune cell subtypes^20^. Different factors such as weaning^21^, viral exposure^22^ or metabolic status^23,24^ influence DNA methylation in peripheral blood mononuclear cells (PBMC). Furthermore, DNA methylation profiles in blood immune cells have been used effectively to accurately detect and characterize age-related biomarkers in various vertebrate species, including pigs^25^.

While the neuro-epigenome of brain regions in piglets has been analysed in terms of stereotypies and enriched maternal environments^26^, the direct effects of environmental enrichment on the DNA methylation of adult PBMC remained unexplored. The aim of this study was therefore to investigate the PBMC methylome of sows of different ages and parity ranks housed under two contrasted environments that could potentially have an impact on the affective states of these animals.

## Materials and methods

### Experimental design and animals

The experiment was conducted at the Chambre Régionale d’Agriculture de Bretagne Experimental Farm (Saint-Nicolas-du-Pélem, France), as detailed in our published study^27,28^. Young nulliparous sows are randomly assigned upon purchase to one of two independent reproductive units, which feature distinct housing environments ^27,28^. They were kept under two separate housing environments: conventional (C, n = 36) and enriched (E, n = 35) reproductive units. In the C unit, the sows were housed in a pen with a concrete slatted floor and minimal enrichment materials, with a minimum space allowance of 2.4 m^2^ per sow. In the E unit, they were accommodated on deep straw bedding with more space per animal (3.5 m^2^). Pregnant sows resulting from crosses between Landrace and Large White breeds, were housed together in groups since their previous weaning until 10 days before their expected farrowing date. Pregnant sows in both units were of different parity ranks (0 to 7 gestations). On gestation day 105 (G105; approximately 10 days before expected farrowing), sows from both units were transferred to individual farrowing pens with slatted floors. They remained there with their litter for four weeks until weaning, after which the sows returned to their previous E or C units. The sows therefore remained in their assigned housing environment throughout their reproductive life. All sows, whatever the housing conditions, were fed individually via an automatic dispensing system, with standardized management and feeding practices across units. All details regarding the animals were reported in Table 1. The trial was carried out over the course of one year, consisting of two replicates: R1 (from November 2021 to February 2022) and R2 (from March to July 2022). In each replicate, we selected 7-8 multiparous sows per environment for blood sampling, totalling 29 animals (C, n = 7 in R1, and n = 8 in R2; and E, n = 7 in R1 and n = 7 in R2). Particular attention was paid to the sows’ genealogy. The number of individuals included in the experimental design is required for methylome analyses, given that modest environmental effects are expected.

**Table 1:** Physiological parameters and genealogy of sows selected for blood sampling. Body weight and age were recorded at the blood sampling time point on G98. A total of 27 different sows were used: 25 different sows (n =13 in C and n =12 in E) were included in the trial; and two additional sows (one in each environment group) were used twice successively into the trial (C_01/C_11 and E_16/E_25). Abbreviations: Rep, replicate assay; C, conventional housing; E, enriched housing. F1 to F16 and M1 to M24: identity of the fathers and mothers of the experimental sows, respectively; DF and DM: different father and different mother, respectively. PHS and MHS: paternal half-sister and maternal half-sister, respectively).

| Library Number | Rep. | Housing | Body weight (kg) | Age (days) | Gestation number | Parity rank | Father | Mother |
| --- | --- | --- | --- | --- | --- | --- | --- | --- |
| C_01 | R1 | C | 297 | 943 | 4 | HP | PHS5_F08 | DM_M02 |
| C_02 | R1 | C | 297 | 945 | 5 | HP | DF_F09 | DM_M03 |
| C_03 | R1 | C | 324 | 803 | 4 | HP | PHS3_F02 | DM_M07 |
| C_04 | R1 | C | 289 | 634 | 3 | LP | DF_F16 | MHS1_M19 |
| C_05 | R1 | C | 283 | 607 | 3 | LP | PHS1_F05 | DM_M09 |
| C_06 | R1 | C | 239 | 496 | 2 | LP | PHS6_F06 | DM_M08 |
| C_07 | R1 | C | 259 | 495 | 2 | LP | PHS4_F13 | DM_M10 |
| C_08 | R2 | C | 217 | 500 | 2 | LP | PHS6_F06 | DM_M23 |
| C_09 | R2 | C | 241 | 502 | 2 | LP | DF_F14 | MHS3_M11 |
| C_10 | R2 | C | 223 | 503 | 2 | LP | PHS6_F06 | MHS1_M19 |
| C_11 | R2 | C | 313 | 1099 | 5 | HP | PHS5_F08 | DM_M02 |
| C_12 | R2 | C | 328 | 951 | 5 | HP | PHS3_F02 | DM_M05 |
| C_13 | R2 | C | 277 | 780 | 4 | HP | DF_F10 | DM_M16 |
| C_14 | R2 | C | 265 | 641 | 3 | LP | PHS7_F12 | DM_M22 |
| C_15 | R2 | C | 297 | 642 | 3 | LP | PHS7_F12 | DM_M21 |
| E_16 | R1 | E | 323 | 944 | 5 | HP | PHS5_F08 | DM_M06 |
| E_17 | R1 | E | 273 | 830 | 4 | HP | DF_F01 | DM_M24 |
| E_18 | R1 | E | 310 | 802 | 4 | HP | DF_F04 | DM_M18 |
| E_19 | R1 | E | 295 | 804 | 4 | HP | PHS2_F03 | DM_M04 |
| E_20 | R1 | E | 299 | 636 | 3 | LP | PHS1_F05 | DM_M17 |
| E_21 | R1 | E | 240 | 494 | 2 | LP | PHS6_F06 | MHS2_M01 |
| E_22 | R1 | E | 240 | 494 | 2 | LP | PHS4_F13 | DM_M12 |
| E_23 | R2 | E | 211 | 503 | 2 | LP | PHS4_F13 | DM_M14 |
| E_24 | R2 | E | 232 | 501 | 2 | LP | DF_F11 | DM_M20 |
| E_25 | R2 | E | 330 | 1092 | 6 | HP | PHS5_F8 | DM_M06 |
| E_26 | R2 | E | 336 | 951 | 5 | HP | PHS2_F03 | DM_M13 |
| E_27 | R2 | E | 247 | 783 | 4 | HP | PHS1_F05 | MHS3_M11 |
| E_28 | R2 | E | 305 | 782 | 4 | HP | DF_F15 | MHS2_M01 |
| E_29 | R2 | E | 243 | 642 | 3 | LP | DF_F07 | DM_M15 |

### Blood sampling, PBMC preparation and genomic DNA preparation

The experimental timeline is summarized in Figure 1A. Blood samples (10 mL blood collection tubes at a final concentration of 1.8 mg EDTA (Ethylene Diamine Tetra Acetic acid) per mL of blood; three tubes collected per sow) were collected by an experienced veterinarian from the anterior vena cava at 98^th^ day of the gestation period (G98, before transfer to individual farrowing pens) and from the mammary vein at the beginning of lactation (12^th^ day; L12), all from the same animals. During experimental replicate R1, sows from enriched housing group were blood sampled first, and inversely, during experimental replicate R2, sows from conventional housing were collected first. These samples were transported in a thermo box with refrigerated rolls and arrived at the laboratory within three hours of the sampling procedure. Each blood sample was notified with sow identification number: a 6-digit number totally independent of age, parity or experimental treatment of the animals, enabling the samples to be processed blindly. Blood formula was immediately determined. Total numbers and percentages of lymphocytes, monocytes and neutrophils were determined with an MS-9® hematology automatic cell counter (Melet Schloesing laboratories, Osny, France; see all data in^27,28^). A part of blood samples was used for the isolation of peripheral blood mononuclear cells (PBMC) according to a standard protocol^29^. Aliquots of 2 million cells were prepared with 0.2 mL lysis buffer (SLB; 10 mM Tris-HCl, pH 7.5, 10 mM EDTA, 10 mM NaCl, 0.2% SDS), thrown into liquid nitrogen for immediate freezing, and then conserved at −80°C until use.

**Figure 1.**
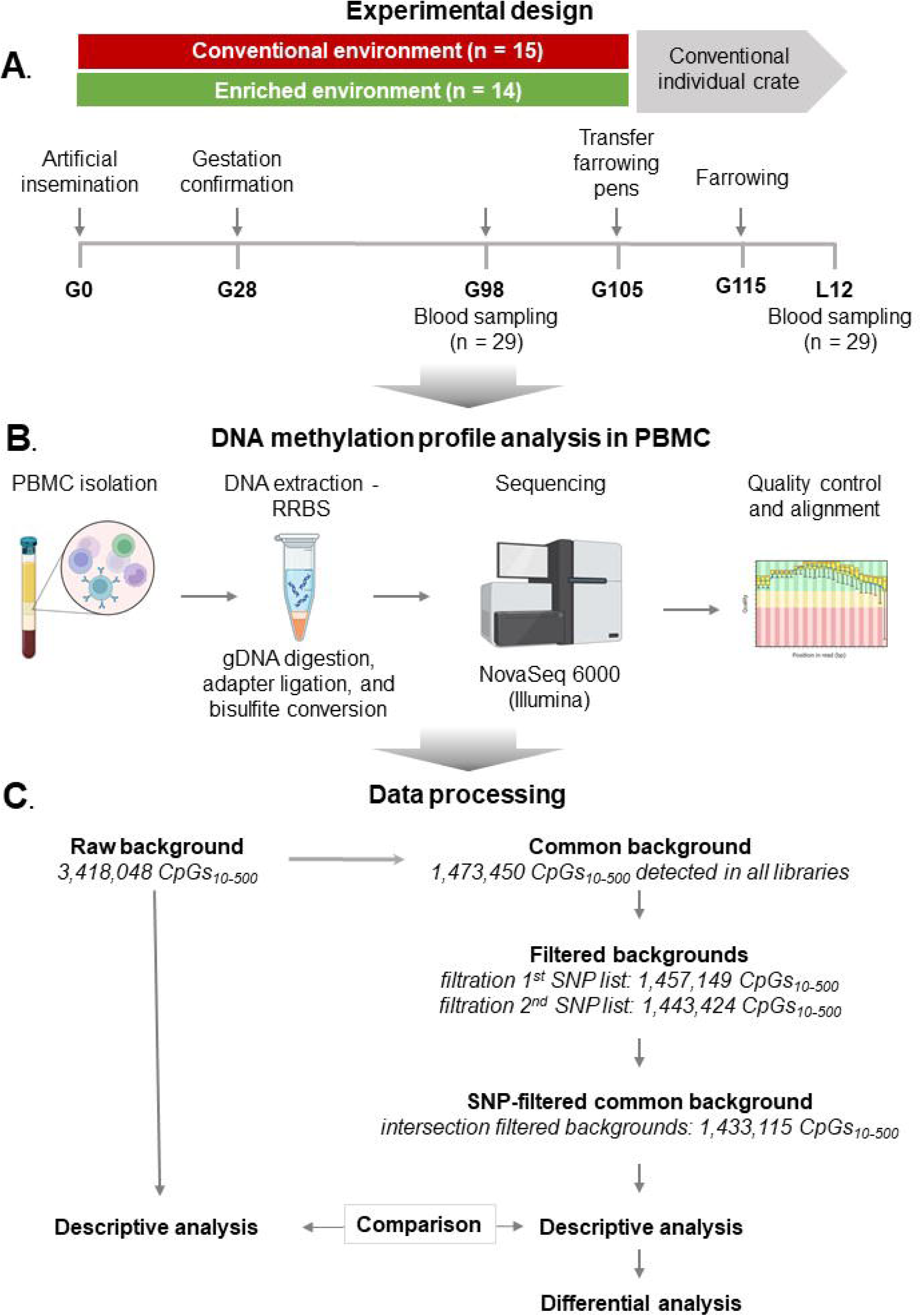
Flowchart and study overview. (A) Schematic outline of the experimental design. (B) Schematic overview of steps from PBMC preparation to the quality control of sequencing data. (C) Schematic overview of CpG background generation and analysis. Abbreviations: G, gestation day; L, lactation day; PBMC, peripheral mononuclear cells; RRBS, reduced representation bisulfite sequencing; CpG_10-500_, CpG with a minimum of 10 and maximum of 500 uniquely mapped reads; SNP, single nucleotide polymorphism.

Genomic DNA (gDNA) extraction was performed following a conventional procedure that included: proteinase K digestion (2mg/mL final concentration) in SLB lysis buffer overnight at 55°C, RNA elimination (RNAse incubation; 0.2mg/mL, 37°C 2h) and two phenol/chloroform extractions. After precipitation with 0.2M NaCl in 70% ethanol (overnight at −20°C and centrifugation, 12,000 x g for 30 minutes at 4°C), the gDNA pellet was washed, dried and resuspended in TE buffer (10 mM Tris-HCl, pH 7.5, 0.1mM EDTA). Integrity (electrophoretic migration on agarose gel), purity (Nanodrop analysis; Thermo Fisher Scientific, Les Ulis, France) and concentration (QuBit fluorometer; Thermo Fisher Scientific) were measured for all gDNA (Figure 1B).

### Reduced Representation Bisulfite Sequencing (RRBS) library preparation

In this study, 58 reduced representation bisulfite sequencing (RRBS) libraries were prepared on a semi-automated machine (Genomic STARlet; Hamilton) according to the procedure previously described by Gu et al.^30^ and adapted by Perrier et al.^31^. gDNA (200 ng) was cleaved by MspI enzyme at CCGG sites, generating fragments with CG dinucleotides at both ends. After the end repair step and Illumina adaptor ligation, a size selection of 150-400 bp of DNA fragments (40 - 290 bp genomic DNA fragments + adapters) was performed using SPRIselect magnetic beads (Beckman-Coulter, Villepinte, France). DNA fragments were then converted twice with sodium bisulphite using the EpiTect bisulphite kit (Qiagen, Les Ulis, France) according to the manufacturer’s instructions, and amplified with Pfu Turbo Cx hotstart DNA polymerase (Agilent, Les Ulis, France) using 14 PCR cycles and Illumina index. Aliquots of PCR products were analysed by electrophoresis on a 4-20% precast polyacrylamide TBE gel (Invitrogen; ThermoFisher scientific, Les Ulis, France) and stained with SYBR green to confirm the homogeneous pattern for all libraries. The libraries were sequenced on an Illumina NovaSeq 6000 sequencer to produce a minimum of 30 million 100 bp paired-end reads (Integragen-OncoDNA, Evry, France).

### Data processing

The data processing method is summarized in Figure 1C. Raw reads underwent quality control using FastQC (v0.12.0; https://www.bioinformatics.babraham.ac.uk/projects/fastqc) and subsequently trimmed using Trim Galore! (v0.4.5; https://www.bioinformatics.babraham.ac.uk/projects/trim_galore) to eliminate reads with a Phred score below 20, shorter than 20 nucleotides, and adaptor sequences. Processed reads were then aligned to the *in-silico* bisulphite-converted *Sus scrofa* reference genome (Ensembl: GCF_000003025.6, Sscrofa11.1) using Bismark v0.20.0 in the default mode with Bowtie 1.2.1^32^. On average, 40 million raw reads per library were generated, with mean total mapping rate of 70% and a mean rate of uniquely mapped reads of 53% covering the pig genome. The bisulphite conversion rate was estimated from the unmethylated cytosine added during the end-repair step. The conversion rate reached 100% in nearly all samples. An overview of sequencing parameters for all libraries is presented in Suppl. Table 1, Sheet 1.

### Selection process of a common CpG_10-500_ background filtered for SNP

Only CpGs covered by 10 to 500 uniquely mapped reads (termed CpG_10-500_) were retained, forming the initial raw background of 3,418,048 CpGs_10-500_, which represents approximately 10% of all CpGs in the pig genome. From this set, only CpG_10-500_ with a methylation value calculated for all the libraries from the G98 and L12 datasets were retained, yielding a background of 1,473,450 CpG_10-500_. To avoid the confounding effects of sequence polymorphisms and bisulfite conversion, the CpGs_10-500_ co-localized with a putative sequence polymorphism affecting the C and/or G were filtered out. For this purpose, the common background (1,473,450 CpG_10-500_) was thus subjected to further filtration using two SNP lists to identify and minimize confounding effects of the animals’ genetic background. The first list was obtained from whole genome sequencing data (accession number PRJEB51909) and genotyping calling performed in 36 Large White pigs as described in^33^. The total of 13,408,342 positions of Large White variants were compared to the positions of CpGs highlighted in the analyses to filter out common positions using bedtool v2.27.1^34^. The remaining CpGs (non-SNP CpGs), were retained for subsequent analyses, yielding 1,443,424 CpGs (30,026 CpGs removed). The second SNP list was extracted from the Farm animal Genotype-Tissue Expression (FarmGTEx) consortium that produced a comprehensive public resource of genetic regulatory variants across tissues and cells in pigs (PigGTEx^35^). This list comprised 3,087,268 SNPs documented in three different breeds: Duroc, Landrace, and Large White. It enabled generation of a filtered background of 1,457,149 CpG_10-500_ (16,301 CpGs removed). Finally, only CpG_10-500_ positions that overlapped between the two filtered backgrounds were conserved (VennDiagram R package) and defined as the final filtered common SNP list consisting of 1,433,115 CpG_10-500_.

### Descriptive analysis of the genome-wide methylation of blood cell libraries

The “methylation_extractor” function from Bismark was used to determine the methylation level at each CpG by counting C and T (methylated and unmethylated before bisulfite conversion, respectively). To monitor the genome-wide methylation pattern before and after SNP filtering, non-supervised hierarchical clustering was conducted using the FactoMineR package^36^ on the two matrices of methylation percentages before (3,418,048 CpGs_10-500_) and after (1,433,115 CpG_10-500_) filtering.

The global mean methylation levels (Suppl. Table 1, Sheet 1) for different sow groups were calculated and statistical analyses were performed using a linear mixed-effect model (lmer function of the R software lme4 package, version v. 4.1.2 (R Core Team, 2023)). The linear model included the replicate as a random factor as well as three fixed factors and their two-by-two interactions: the physiological stage (G98 or L12), the parity class (classified in two groups: low-parity (LP; ranks 2-3) and high-parity (HP, ranks 4 and higher) and the housing environment (E or C).

### Identification of differentially methylated cytosines/regions (DMCs/ DMRs)

Each differential methylation analysis between the two groups was conducted using DSS software (Dispersion Shrinkage for Sequencing data, v2.46.0^37^) in the default mode. Differentially methylated cytosines (DMCs) were identified considering a minimum methylation difference between groups of 15%, and the adjusted *p* value ≤ 0.05. This *p* value adjustment was performed according to the Independent Hypothesis Weighting method, using the alpha parameter set at 5% and the average methylation per group as a covariable. Moreover, DMRs (Differentially Methylated Regions) were identified and defined as regions containing at least three DMCs with a distance between each DMC of 100 bp or less. For each DMC, a particular attention was focused on individual methylation values distribution. It was observed either continue distribution (values varying from 0 to 1) or tri-modal distribution values distributed around of 0 (unmethylated), or around of 0.5 (intermediate-methylated) or around of 1 (methylated). The tri-modal distribution suggests an overlapping SNP/CpG, which can thus be misidentified as unmethylated Cs: indeed, C>T SNPs cannot be distinguished from C>T substitutions that are caused by bisulfite conversion and G>A SNPs induces a loss of methylation on C previous G.

### Annotation for genomic elements

The final SNP-filtered common background, DMCs and DMRs were annotated based on their genomic location using reference files obtained from the Ensembl website (ftp://ftp.ensembl.org/pub; release101). They were classified relative to gene features that included exons, introns, transcription start sites (TSS; DMCs located within 100 base pairs up- or downstream of TSS), promoters (DMCs located within −2000 /+100 base pairs of TSS) and transcription termination sites (TTS; DMCs located within 100 base pairs up- or downstream of TTS). A DMC/DMR was considered to belong to a CpG island if there was at least 75% overlap between the site/fragment and the CpG island, and to a shore (up to 2000 base pairs from a CpG island) or shelf (up to 2000 base pairs from a shore). A DMC/DMR was considered as being overlapped by a repetitive element whatever the extent of this overlapping. In order to better characterize the DMRs detected during this study, we reassessed the initially assigned genomic features of a subset of DMRs by intersecting them with functional annotation data (Ensembl Regulation dataset): promoters, enhancers, open chromatin regions, CTCF bindings sites (https://ftp.ensembl.org/pub/release-113/regulation/sus_scrofa/Sscrofa11.1/annotation/Sus_scrofa.Sscrofa11.1.regulatory_features.v113.gff3.gz) and, Epigenetically Modified Accessible Regions (EMARs) (https://ftp.ensembl.org/pub/release-113/regulation/sus_scrofa/Sscrofa11.1/annotation/Sus_scrofa.Sscrofa11.1.EMARs.v113.gff.gz)

### Gene Ontology and KEGG enrichment analysis

The list of genes co-localized with the resulting DMCs were subjected to enrichment analysis with the DAVID bioinformatic database v2022q2^38^, which included Gene Ontology biological process (GO-BP) terminologies and KEGG pathway categories. The whole list of genes co-localized with SNP-filtered common CpG_10-500_ was used as the genome reference. Pathways with at least three genes and p-values <0.05 were considered to be enriched. We also used the gene functional annotations from GeneCards and Ensembl Regulation databases.

### Statistical analysis of blood cell counts and global DNA methylation level

Linear mixed-effect models, were applied for statistical analysis using lmer function of the lme4 package (under R version v. 4.1.2). In these models, the sampling day as fixed effect and sow as random effect were included in order to consider the repeated measures on each sow. Factor effects were considered significant if the p-value was < 0.05.

## Results

### Blood leucocytes proportion

Blood samples were collected at two time-points (G98 and L12) and blood formula was determined. No significant differences in blood cell sub-population numbers were observed between sows housed in the two environments (p > 0.05). Higher lymphocyte levels were seen in the low-parity (LP) group compared to the high-parity (HP) group of sows (see below, p < 0.05), and significant differences between sampling dates (G98 vs. L12) were observed for monocyte and neutrophil counts (p < 0.001 for both, Table 2). From the blood samples, PBMC preparations were performed and used to determine DNA methylation profiles. Although the lymphocytes- and monocytes-count variations are limited, it may to be considered subsequently for methylation data interpretation.

**Table 2.** Effect of environmental enrichment blood cell numbers. Values are presented in raw values (mean ± sem). Abbreviations: C, conventional; E, enriched; HP, high-parity; LP, low-parity; H, housing; P, parity; D, dates; G, gestation day; L, lactation day.

|  | HP |  | LP |  | *p-value* |  |  |
| --- | --- | --- | --- | --- | --- | --- | --- |
|  | C<br>(n = 6) | E<br>(n = 8) | C<br>(n = 9) | E<br>(n = 6) | H | P | D |
| Lymphocytes<br>(10<sup>3</sup>/μL) | 4.4 ± 0.1 | 4.0 ± 0.1 | 4.7 ± 0.1 | 4.9 ± 0.2 | 0.5 | 0.02 | 0.1 |
| Monocytes<br>(10<sup>3</sup>/μL) | 1.4 ± 0.1 | 1.2 ± 0.1 | 1.3 ± 0.07 | 1.4 ± 0.1 | 0.8 | 0.8 | < 0.001 |
| Neutrophils<br>(10<sup>3</sup>/μL) | 6.0 ± 1.3 | 4.4 ± 0.3 | 4.9 ± 0.3 | 5.2 ± 0.7 | 0.3 | 0.9 | < 0.001 |

### Description of high inter-individual variability in DNA methylation profiles

DNA methylation profiles were initially explored using the initial raw data (3,418,048 CpGs_10-500_ before filtering; Suppl. Figure 1) and the SNP-filtered background (1,433,115 CpG_10-500_. (Figure 2A). Both dendrograms displayed quite similar overall structure. Non-supervised hierarchical clustering revealed one cluster with 30 libraries and the other with 28. As expected, for all individual, libraries at G98 and L12 clustered together. Moreover, a distinct cluster of four samples (two physiological stages x two replicates) was observed for each two sows used in both replicates. The SNP-filtering reduced distance scale indicated that the suppression of SNP-overlapping CpGs contributed to a reduction in inter-individual variations. Therefore, the following analyses are only presented with methylation data for the SNP-filtered background.

**Figure 2.**
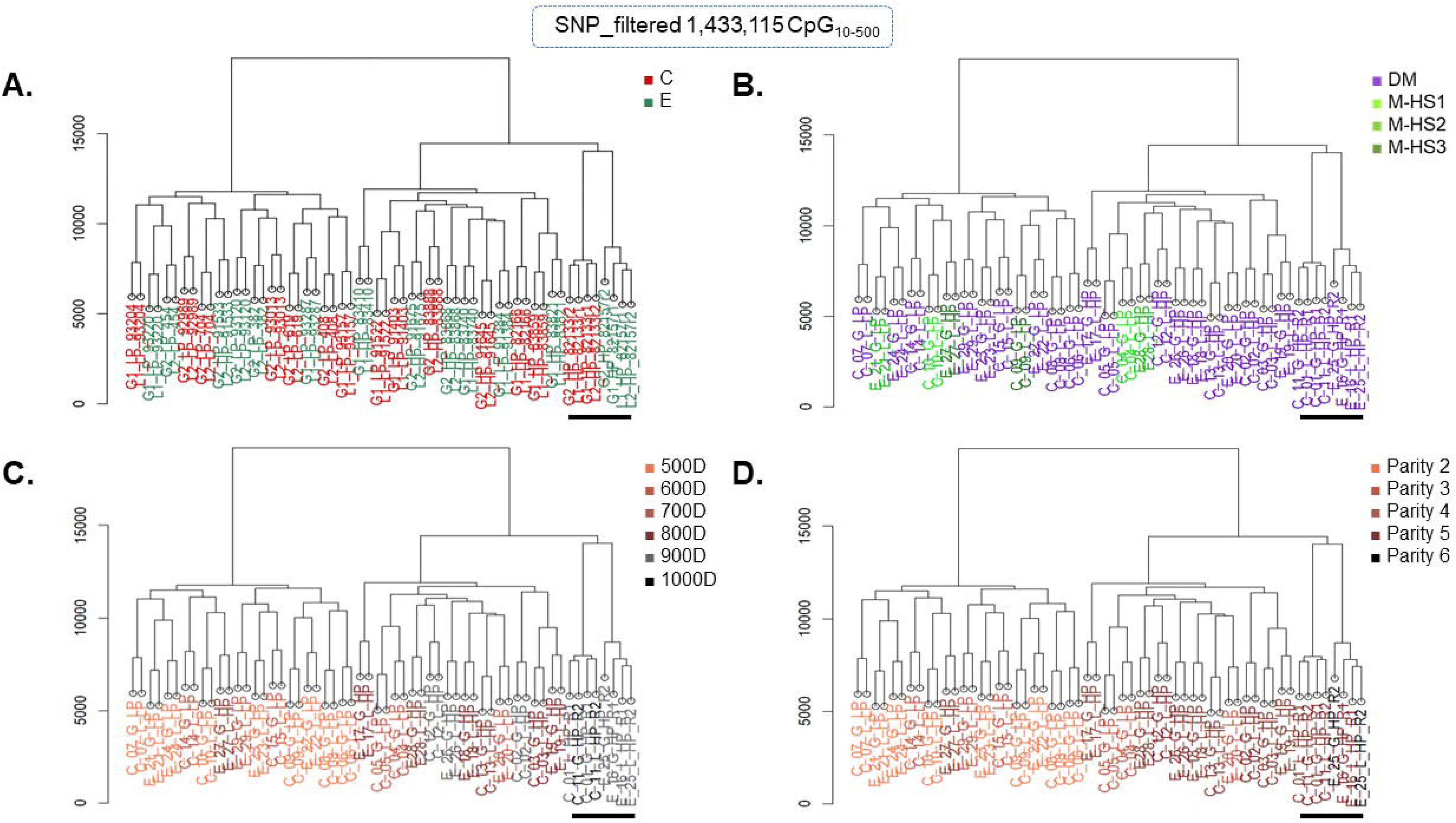
Hierarchical non-supervised clustering using a matrix obtained after SNP-filtering and including the methylation level of 1,433,115 CpG_10-500_. The library names are coloured according to (**A**) housing environments, (**B**) maternal genealogy (**C**). sow age range and (**D**) sow parity rank. Abbreviations: C, conventional; housing E, enriched housing; DM, different mother; MHS, maternal half-sisters; D, days of age; as described in Table 1. Black bar underlines the libraries produced from the sows included in both replicates (8 different libraries).

Surprisingly, clustering did not separate samples based on housing environment (C vs. E, Figure 2A). To better understand the factors influencing hierarchical clustering, we first compared the two replicates of the experimental design (R1 vs. R2) and found no discernible clustering pattern (Suppl. Figure 2A), suggesting an absence of significant technical biases or seasonal effects.

### Sows’ genealogy was not the illustrative factor of clustering

When categorized by maternal origin, we identified 19 sows with no half-sisters (38 libraries from G98 and L12) and six sows with at least one half-sister (12 libraries from G98 and L12). Maternal genealogy did not significantly influence the two branches of the dendrogram (Figure 2B). We then examined paternal origins, categorizing sows into nine with no half-sisters (18 libraries from G98 and L12) and 16 with at least one half-sister (32 libraries from G98 and L12; Suppl. Figure 2B). Similarly, paternal origins did not significantly influence the clustering, before or after filtering.

### Age and parity class clearly influenced the individual DNA methylation variability

The age of sows at G98 ranged from 494 to 1,099 days. Hierarchical clustering revealed a clearer separation based on age (Figure 2C): the first branch (n = 26 libraries) included sows aged 400 to 599 days, while the second branch (n = 32 libraries) included sows aged 600 to 1,099 days. A similar pattern emerged when clustering was analysed by parity class (Figure 2D): the first branch consisted of sows with 2-3 parities, while the second branch included sows with 4 or more parities. Only four pairs of libraries did not cluster as expected: E_27 from HP group was clustered with LP group and conversely, C_04, C_05, E_20 from LP group - with HP group. Nevertheless, no clear explanation has been found.

In light of these results, a new categorization of the sows based on parity class was made: those in their second or third reproductive cycle (547 ± 73 days old at G98 sampling) formed the low-parity group (LP, n = 15), while those in their fourth or subsequent cycle (860 ± 163 days old at G98 sampling) formed the high-parity group (HP, n = 14).

We then investigated the effects of parity (HP vs. LP), physiological state (G98 vs. L12), and housing environment (E vs. C) on global DNA methylation percentages (Suppl. Table 2). The mean global DNA methylation of PBMCs, calculated across all filtered CpGs (1,433,115 CpGs), was 54.93% ± 0.53. No significant differences in global methylation were observed between sows from the E and C environments (p > 0.05, Figure 3A), although limited, but significant differences were found between parity groups (Figure 3B) and physiological states (Figure 3C; p < 0.01 for both).

**Figure 3.**
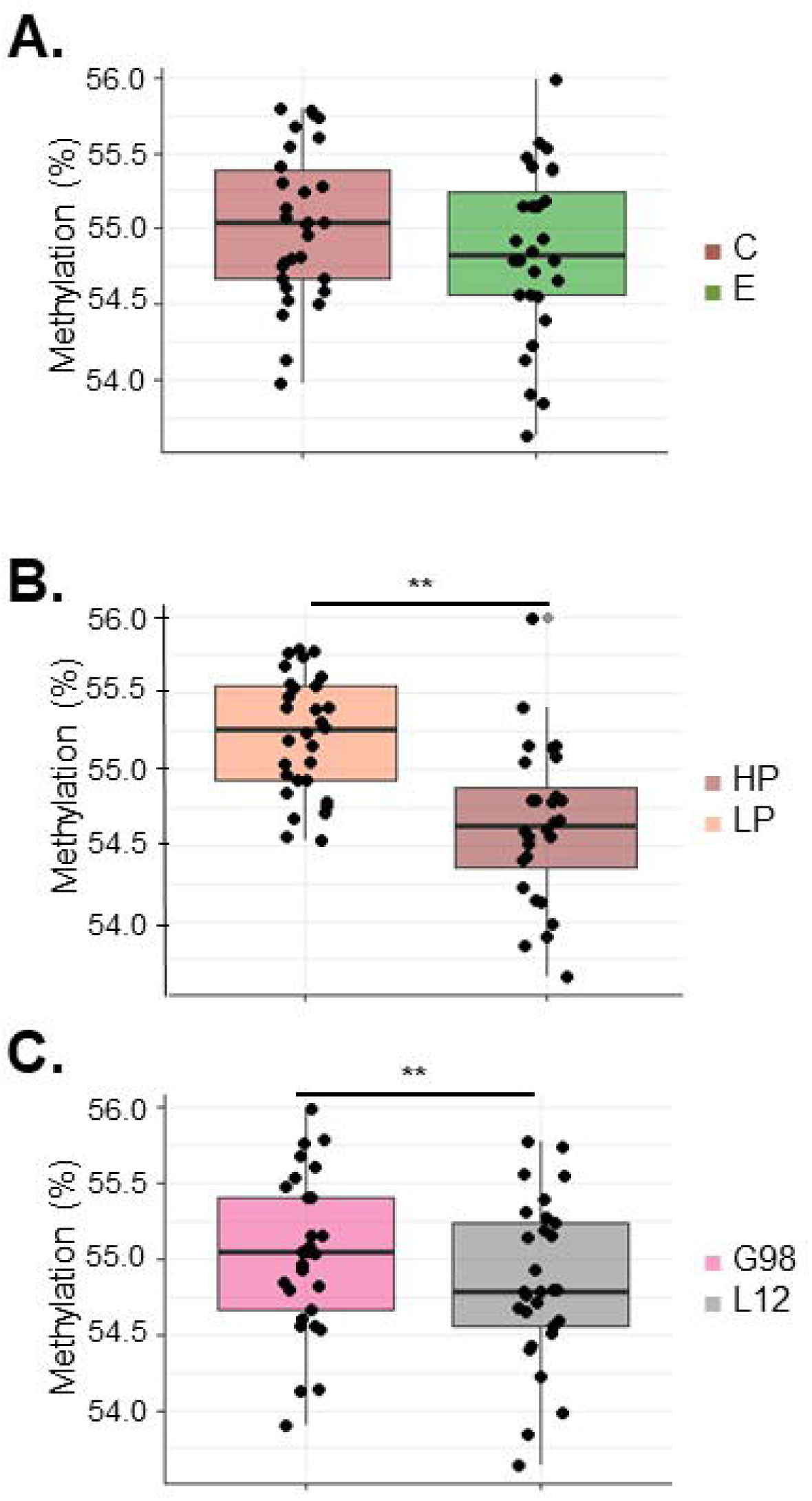
Effects of different factors on DNA global methylation percentages in samples. (A) Comparison of conventional (C) and enriched (E) groups, (B) of low-parity (LP) and high-parity (HP) groups, and (C) gestation day 98 (G98) and lactation day 12 (L12) physiological states. ** P <0.01.

### Identification of differentially methylated cytosines (DMCs) confirmed the strong influence of parity

When comparing HP and LP sows at day G98, we identified 1,358 DMCs, displaying a global reduction in methylation observed in HP sows (1,011 less and 347 more methylated; Suppl. Table 1, Sheet 2). The chromosomal distribution of DMCs (Figure 4) indicated that chromosomes 6, 1, 12, 2, and 3 contained the highest number of DMCs, independently of their size. Chromosome 12, in particular, displayed a high density of DMCs, probably due its higher GC content (47.5% versus a mean of 42.1 ± 2.0% for all chromosomes). Except for chromosomes 10, 18, and X, DMCs were enriched in the terminal regions of most chromosomes. This pattern may partly have resulted from the RRBS method, that targets CpG-rich regions often found at chromosome ends (Suppl. Figure 3A).

**Figure 4.**
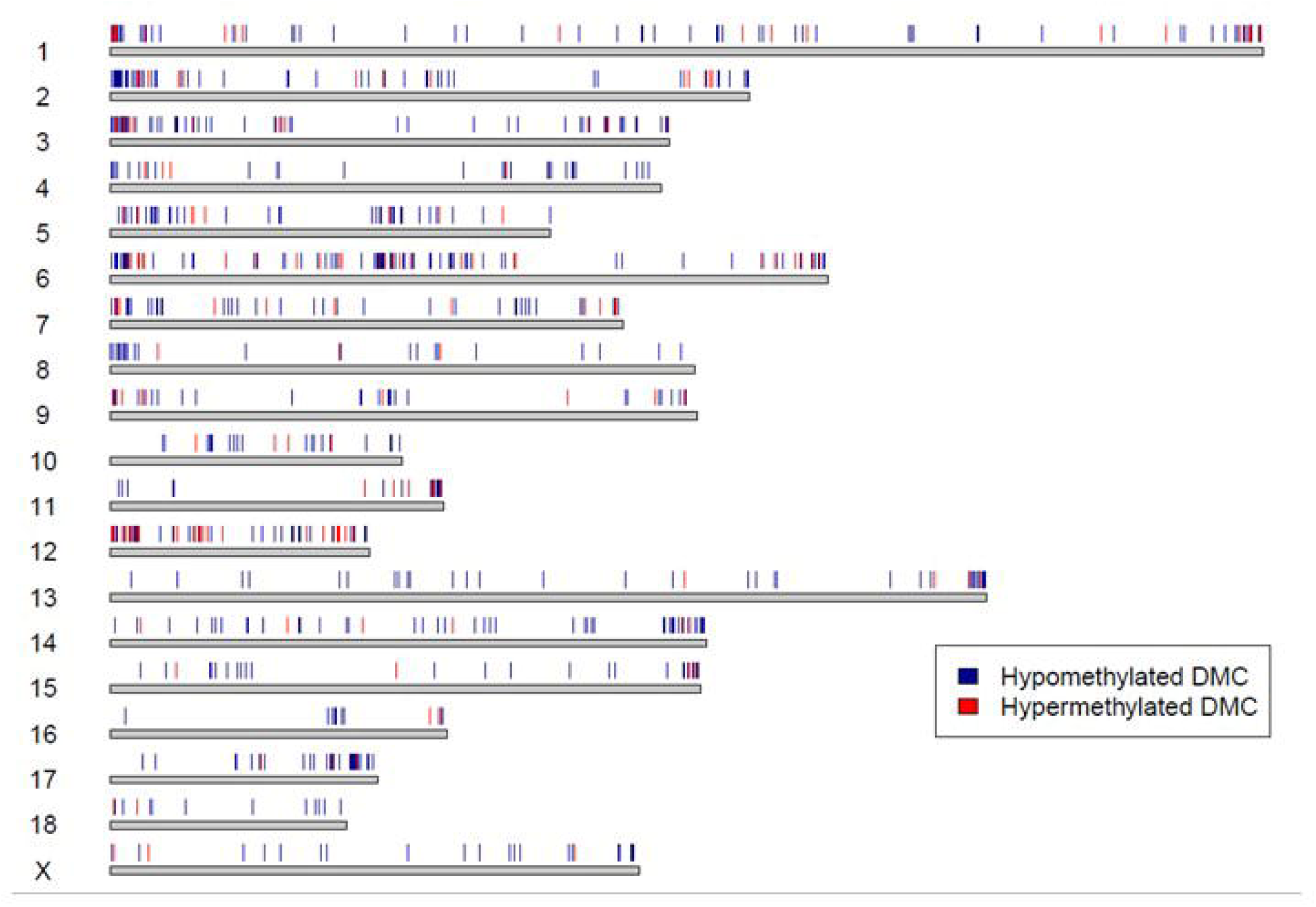
Genome-wide distribution of parity related -DMCs, identified at G98. Pig 18 autosomes and X chromosomes are represented by horizontal bars according to their size (https://www.ncbi.nlm.nih.gov/datasets/genome/GCF_000003025.6/). DMCs identified between low and high parity on G98 are represented by coloured thin lines: blue lines for less methylated DMCs and red lines for more methylated DMCs in the HP group.

The analysis of HP and LP sows at day L12 revealed 680 DMCs (552 less methylated and 128 more methylated in HP sows; Suppl. Table 1, Sheet 3). While the number of DMCs was lower at L12 than at G98, a similar trend of less methylation was observed in HP sows (81.2% of DMCs), with a comparable genomic and chromosomal distribution. However, DMC enrichment at the chromosome termini was less pronounced at L12 (Suppl. Figure 3B).

While a small portion of the DMCs were shared between the two time-points (252 DMCs, accounting for 18.5% of G98 DMCs and 37% of L12 DMCs), a similar genomic distribution of DMCs was observed at both stages (Table 2). A significant portion of DMCs was located in intergenic regions (30.3% and 32.3% at G98 and L12, respectively). Among genic regions, introns contained the most DMCs (40.4% and 39.4%), followed by exons (9.4% and 7.5%) and 5’UTRs (8.8% and 8.6%). Most DMCs were located in CpG island (CGi) desert (54.8% and 56.7%), with the remainder distributed across CGi, CGi shores and CGi shelves. The functional analysis revealed that the 698 genes targeted by DMCs at G98 were involved in 15 enriched biological processes (BPs), while 390 genes at L12 were involved in 10 enriched BPs (Table 3).

**Table 3.** Genomic distribution of differentially methylated cytosines (DMCs) between high-(HP) and low-parity (LP) sows on gestation day 98 (G98) and lactation day 12 (L12); ^1^ Numbers and calculated percentages are indicated. Percentages were calculated relative to the total numbers of DMCs at each time point. ^2^As a reference, the genomic distribution of 1,433,115 CpG_10-500_ (filtered and common CpG_10-500_) was determined. The box colour shows whether a preferentially genomic distribution of DMCs was observed compared to that of filtered and common CpG_10-500_ (green = increase of % and red = decrease of %).

|  | DMCs<br>G98<sup>1</sup> | DMCs<br>L12<sup>1</sup> | CpGs<sub>10-500</sub><br>background<sup>2</sup> |
| --- | --- | --- | --- |
| **CpG in gene bodies versus intergenic regions** |  |  |  |
| Intergenic | 412<br>(30.3%) | 220<br>32.3 % | 284,740<br>19.8 % |
| Exon | 128<br>(9.4%) | 51<br>(7.5 %) | 228,050<br>(15.9 %) |
| Intron | 569<br>(40.4%) | 268<br>(39.4 %) | 479,396<br>(33.4 %) |
| Promoter | 68<br>(5 %) | 46<br>(6.7 %) | 134,874<br>(9.4 %) |
| TSS | 15<br>(1.1 %) | 3<br>(0.04%) | 83,225<br>(5.8 %) |
| TTS | 1<br>(0.07%) | 0<br>(0%) | 5,562<br>(0.3 %) |
| 3'UTR | 45<br>(3.3 %) | 33<br>(4.8 %) | 51,146<br>(3.5 %) |
| 5'UTR | 120<br>(8.8 %) | 59<br>(8.6 %) | 166,120<br>(11.5 %) |
| **CpG associated with CpG islands** |  |  |  |
| CpG desert | 745<br>(54.8 %) | 386<br>(56.7 %) | 494,291<br>(34.4 %) |
| CpG Island | 246<br>(18.1 %) | 128<br>(18.8 %) | 701,098<br>(48.9 %) |
| CGi shelves | 149<br>(10.9 %) | 50<br>(7.3 %) | 86,408<br>(6%) |
| CGi shores | 218<br>(16 %) | 116<br>(17 %) | 151,316<br>(10.5 %) |
| **Overlapping with repeat sequences** |  |  |  |
| Without overlapping | 984<br>(72.5 %) | 468<br>(68.8 %) | 1,123,127<br>(78.3 %) |
| With overlapping | 374<br>(27.5%) | 212<br>(31.2 %) | 309,986<br>(21.6 %) |

**Table 4.**
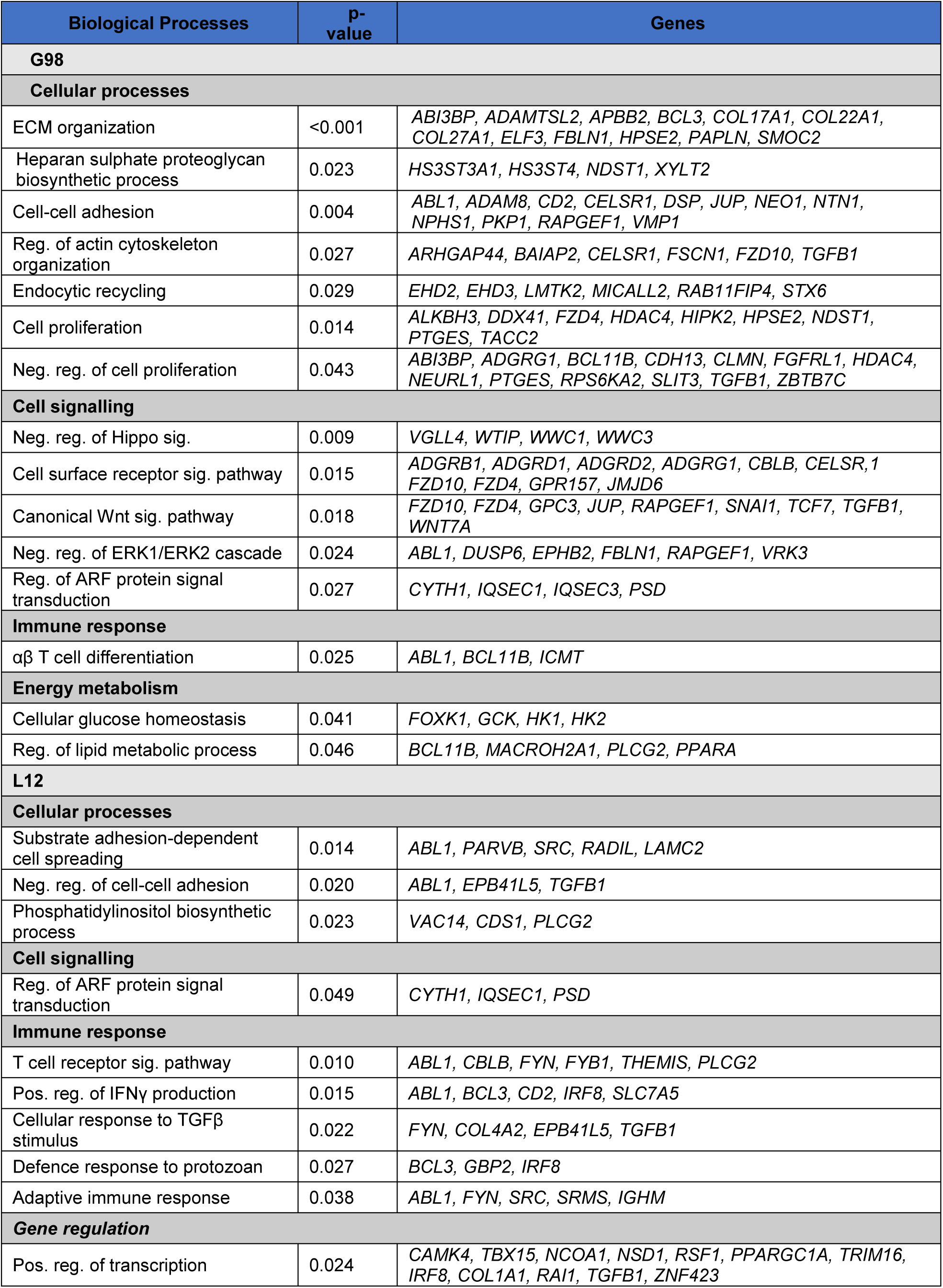
Functional enrichment of DMC-associated genes linked with parity class.

Further analysis looked at the distribution of methylation values for the 252 DMCs common to. Both physiological stages. Among them, 51.6% of these shared DMCs exhibited a tri-modal methylation distribution (around of 0 - unmethylated; around of 0.5 - intermediate-methylated; around of 1 - methylated), likely influenced by genetic factors, while 44% displayed a continuous distribution, indicative of a potential epigenetic effect. A small percentage (4.4%) had an unclear pattern (Suppl. data 1.pdf).

### Differentially methylated regions (DMRs) also highlight the impact of parity rank on DNA methylation profiles

At G98, 60 DMRs were identified (58% of them being less methylated and 42% more methylated in HP), spanning 52 unique genes including 9 uncharacterized genes (Suppl. Table 1, Sheet 4 and Suppl. Table 3). Most DMRs were found in intronic (30%) and exonic (16.7%) regions, followed by promoter-associated regions (18.3% in 5’UTRs, 8.3% in promoters, and 1.7% in TSS), 3’UTRs (5%) and ± 10kb of gene (11.6%); only 8.3% of DMRs were localized in intergenic regions. At L12, only 24 DMRs were identified, evenly split between loss and gain of methylation in HP, encompassing 23 unique genes including 7 uncharacterized genes and similarly distributed on gene features than G98-DMRs (Suppl. Table 1, Sheet 4 and Suppl. Table 3).

Only a few DMRs displayed a tri-modal methylation distribution (3 at G98 and two at L12 Suppl. data 2.pdf and Suppl. data 3.pdf). In particular, the *MYLK* DMR including seven DMCs, is in 5’UTR of gene and co-localized with a CGi shelf (Figure 5A). A tri-modal distribution of methylation levels was observed: among the LP group, most of them had a methylation level around 50%, while the most of HP group had methylation levels around 100%. Few individuals displayed a methylation level at 0% (Figure 5B). The tri-modal pattern was highly correlated between G98 and L12 in both LP (Figure 5C, R² = 0.84) and HP (Figure 5D, R² = 0.93) animals. These results illustrated genetics-epigenetics interactions at the *MYLK* 5’UTR region.

**Figure 5.**
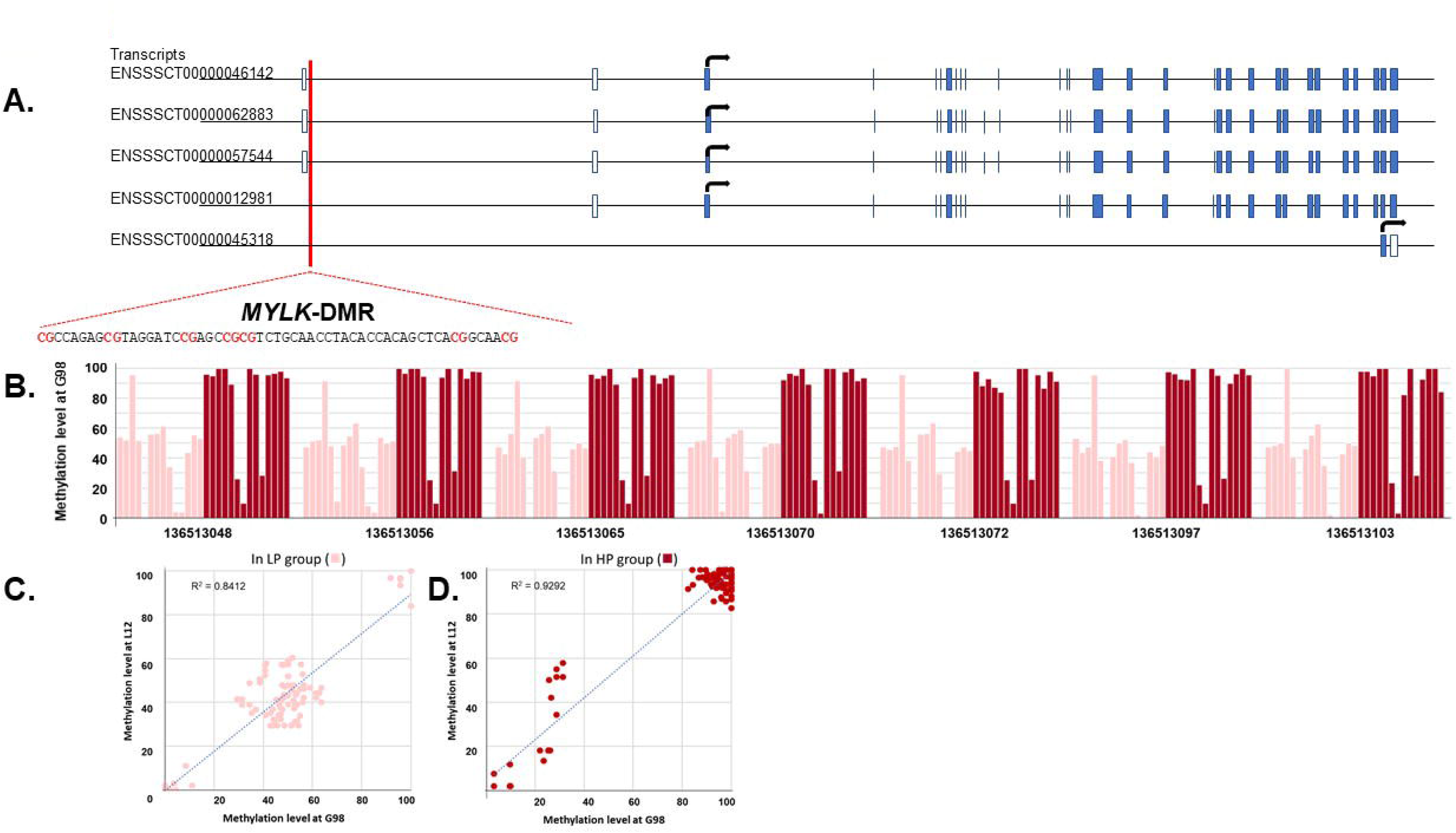
*MYLK*-DMR. (**A**) Chromosomic position, gene organization and DMR position and sequence ((in red, CG positions identified as DMC). (**B**) Methylation levels of DMCs (named by the start genomic position) for each individual from LP and HP groups at G98; Black arrow indicates TSS. (C) Correlation between methylation levels at G98 and L12 in the LP group. (D) Correlation between methylation levels at G98 and L12 in the HP group.

Nevertheless, 55 parities associated DMRs (44 at G98 and 15 at L12 including 4 in common; Suppl. Table 3) exhibited a continuous methylation distribution, suggesting an epigenetic effect of parity class. Although the methylation levels at G98 and L12 were not significantly correlated, the parity-related differential methylation pattern was preserved at both stages for *CCND3* (Figure 6A-D), *HDAC4*, and *SGK1* (Suppl. Figures 4). We re-evaluated the genomic features of the 55 DMRs (Suppl. Table 3) by intersecting them with functional annotation data (Ensembl Regulation dataset). Of these, 15 DMRs (25.86%) co-localized with enhancers, 12 DMRs (20.68%) co-localized with epigenetically modified accessible regions/open chromatin regions and, one co-localized with CTCF binding site; highlighting their potential regulatory role in gene expression control. Together the results from gene feature annotation combined to functional annotation, highlighted the fundamental regulatory role of DNA methylation changes observed with parity rank. More remarkably, DMRs-associated with DNA methylation pattern altered in HP sows encompassed genes which are putatively associated with immune response as schematically presented (Figure 7). Some DMRs presented a loss of methylation in HP sows on G98, encompassing genes such as *CD2* (in promoter)*, CAMK4* (in 5’UTR)*, SECTM1* (in 5’UTR - co-localized with an enhancer), *UMODL1* (in upstream region) and *CMIP* (in intron – co-localized epigenetically modified accessible region). Some DMRs displayed a gain of methylation on G98, such as in intron of *SEC14L1* (co-localized with an enhancer), intron of *SKI* (co-localized epigenetically modified accessible region), 5’UTR of *PACS1* and intron of *SGK1-DMR* (co-localized with an enhancer and CGi-shore). Some DMRs were only observed at L12: they co-localized with intron of genes such as *CD5* and *TNFRSF1B* and displayed a loss of methylation. One of the two *SGK1*-DMRs as well as the *CCND3-*DMR (colocalized in 5’UTR) were found at both time-points and displayed an increase of methylation level in HP sows. Four DMRs were associated with genes involved in epigenetic processes: *DNMT3A* (more methylated in intron – co-localized with an enhancer) and *KDM8* (more methylation in exon) at G98, *SIRT2* (less methylated in promoter) at L12 and *HDAC4-*DMR (less methylated in intron – co-localized with an enhancer) at both time-points. Other DMRs were associated with protein coding genes with functional roles in cell interactions with the extracellular matrix (*EPB41L5, ICAM2, TNNT3, UMODL* at G98), cell adhesion (*AMIGO3, THBS2* at L12), signalling cascades *(DUSP6, GPR68, GRK4, RAPGEF1* at G98 and *RASL10B, GRAP2* at L12), transcription, mRNA splicing and translation control (*SCAF8*, *UNK*, *CELF5*, *ZNF205*, *POLA2*, *FLYWCH2, TCF20, SKI*), and ubiquitin-specific processes (*USP36*, *NSMCE1*, *KLHL36*, *RCHY1*). Taken together, these results point to that the reproductive life, itself evolving with the age of the individual, induces changes in immune methylation profiles, which could play a central role in controlling the immune response.

**Figure 6:**
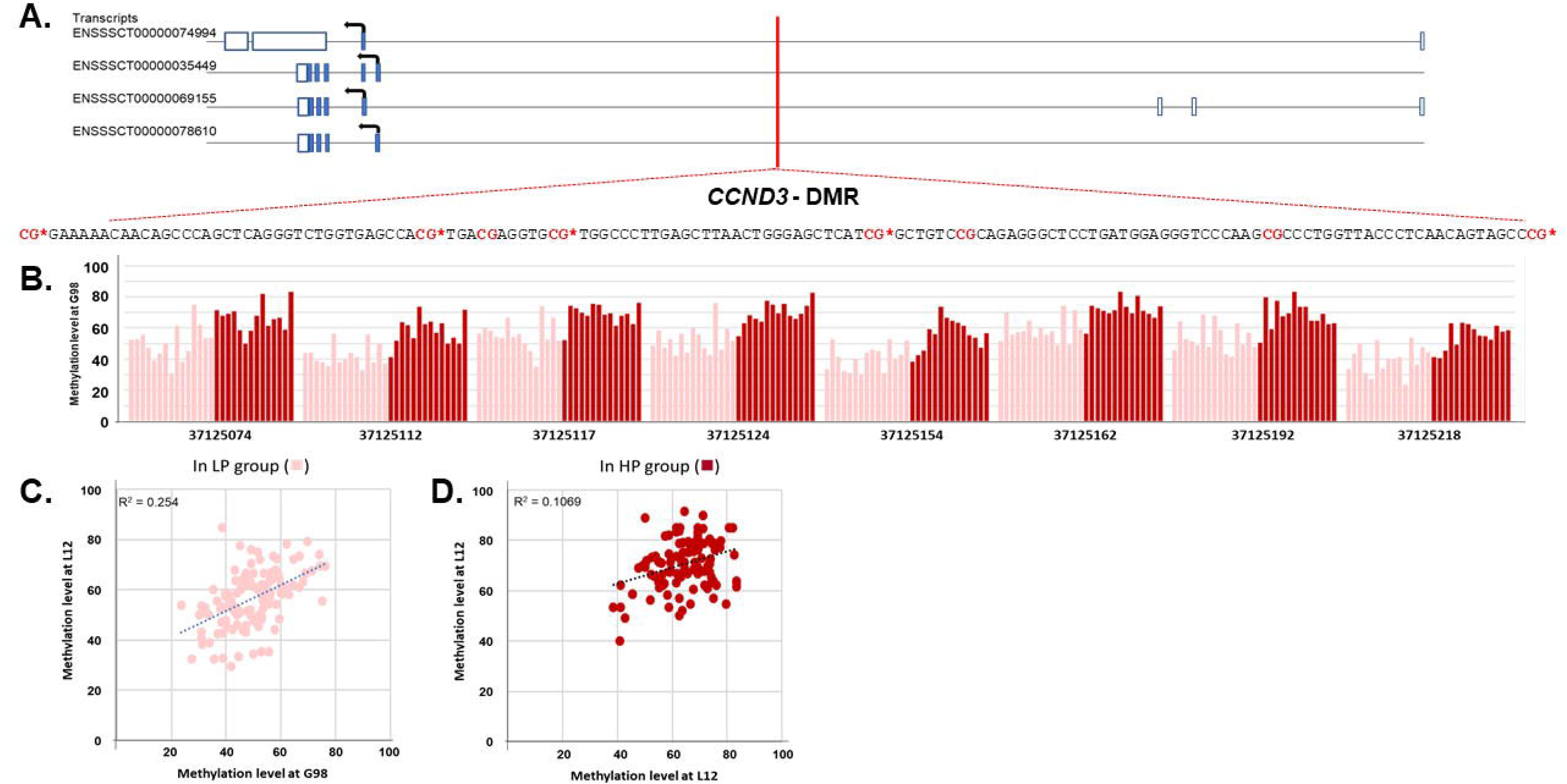
*CCND3*-DMR. (**A**) Chromosomic position, gene organization and DMR position and sequence (in red, CG positions, * indicates them identified as DMC). (**B**) Methylation levels of DMCs (named by the start genomic position) for each individual from LP and HP groups at G98; Black arrow indicates TSS. (C) Correlation between methylation levels at G98 and L12 in the LP group. (D) Correlation between methylation levels at G98 and L12 in the HP group.

**Figure 7.**
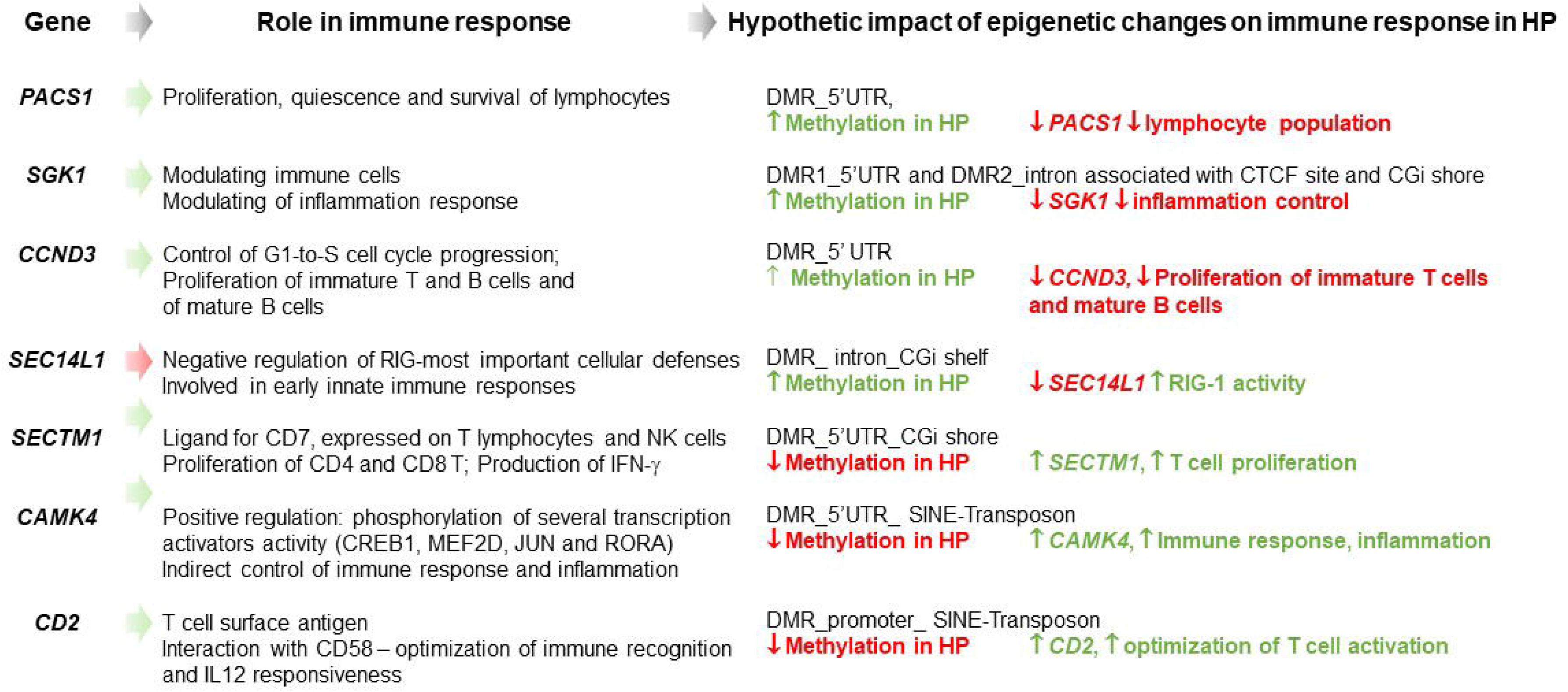
Focus on genes, affected by epigenetic changes induced by parity class and hypothetic impact on immune functions. Only related parity-DMRs colocalized in promoter and/or upstream regulatory region of gene were selected: thus, seven genes were presented. They encode proteins with role in immunity based on information from GeneCards, the Human Gene database (https://www.genecards.org/).

### The DNA methylation changes in response to housing enrichment depended on sow’s parity

Given the significant impact of parity class observed on DNA methylation profiles, we investigated the effects of environmental enrichment (E vs. C) separately for each parity group, first in LP sows (E = 6, C = 9) and then in HP sows (E = 8, C = 6). At G98, we identified 60 DMCs impacted by the housing environment in LP sows (25 less - and 35 more methylated in E) and 42 in HP sows (23 less and 19 more methylated in E). At L12, 35 housing-associated DMCs were seen in LP animals (18 less - and 17 more methylated in E) and 81 in HP animals (46 less - and 35 more methylated in E). No DMCs were shared between the two parity groups whatever their physiological stages, suggesting that the reproductive life of animals may influence the epigenetic response to environmental enrichment (Suppl. Table 1, Sheets 5 and 6). No DMRs were identified as a response to environmental enrichment.

As previously, the distribution of methylation profiles for each housing-associated DMC was assessed and categorized as either tri-modal or continuous (Suppl. Table 1, Sheets 5-6; Suppl data 4-7). At G98, whatever the parity group, around 53% of DMCs were classified as continuous suggesting an epigenetic effect. Around 42% were classified as tri-modal suggesting a genetic effect, while 5% exhibited an undetermined status. At L12, only 44 % of DMCs were classified as continuous and 66% tri-modal and/or with undetermined status. The genomic location of the housing-associated DMCs (Table 5) was more often in CpG desert or intergenic regions, and, when found in genic regions, it was mainly situated in intronic areas. Due to the limited number of DMCs identified, pathway enrichment analysis was not feasible. Nevertheless, we were able to classify manually the annotated DMC-associated genes (Suppl. Table 4). For example, some DMCs co-localized with genes involved in immune system and inflammatory response. Two DMCs in TSS-promoter of *NOS2* (Figure 8A), displayed a decrease of methylation in E-LP-G98 group, one DMC upstream to *CHID1* gene was co-localized with an enhancer and exhibited a gain of methylation in E-HP-L12 (Figure 8G). Some DMCs colocalized genes involved in cell cycle control and cell adhesion/migration: one DMC in intron of *MALD1L1* displayed a loss of methylation in E_LP_G98 (Figure 8B) such as one DMC in exon of *LINGO 3* was co-localized with a CGi in E-HP-L12 (Figure 8E). Interestingly, some DMCs also colocalized with genes involved in neuronal functions: in E-LP-G98 group, two DMCs in intron of *SHANK2* gene colocalized with a CpG island and displayed a gain of methylation (Figure 8C), while three DMCs in intron of *SRGAP2* gene colocalized with a type I transposon/SINE and displayed a loss of methylation with enriched environment (Figure 8D). In E-HP-L12, one DMC in intron of *SORCS2* co-localized with CGi shelves (Figure 8F) and one DMC colocalized in upstream region of *SLC1A3* gene (Figure 8H) exhibited a loss of methylation with enriched environment. Interestingly, two different DMCs (related to *CHILD1* and *LINGO 3*) were colocalized with regulatory elements, suggesting a putative role of methylation changes in transcription control.

**Figure 8.**
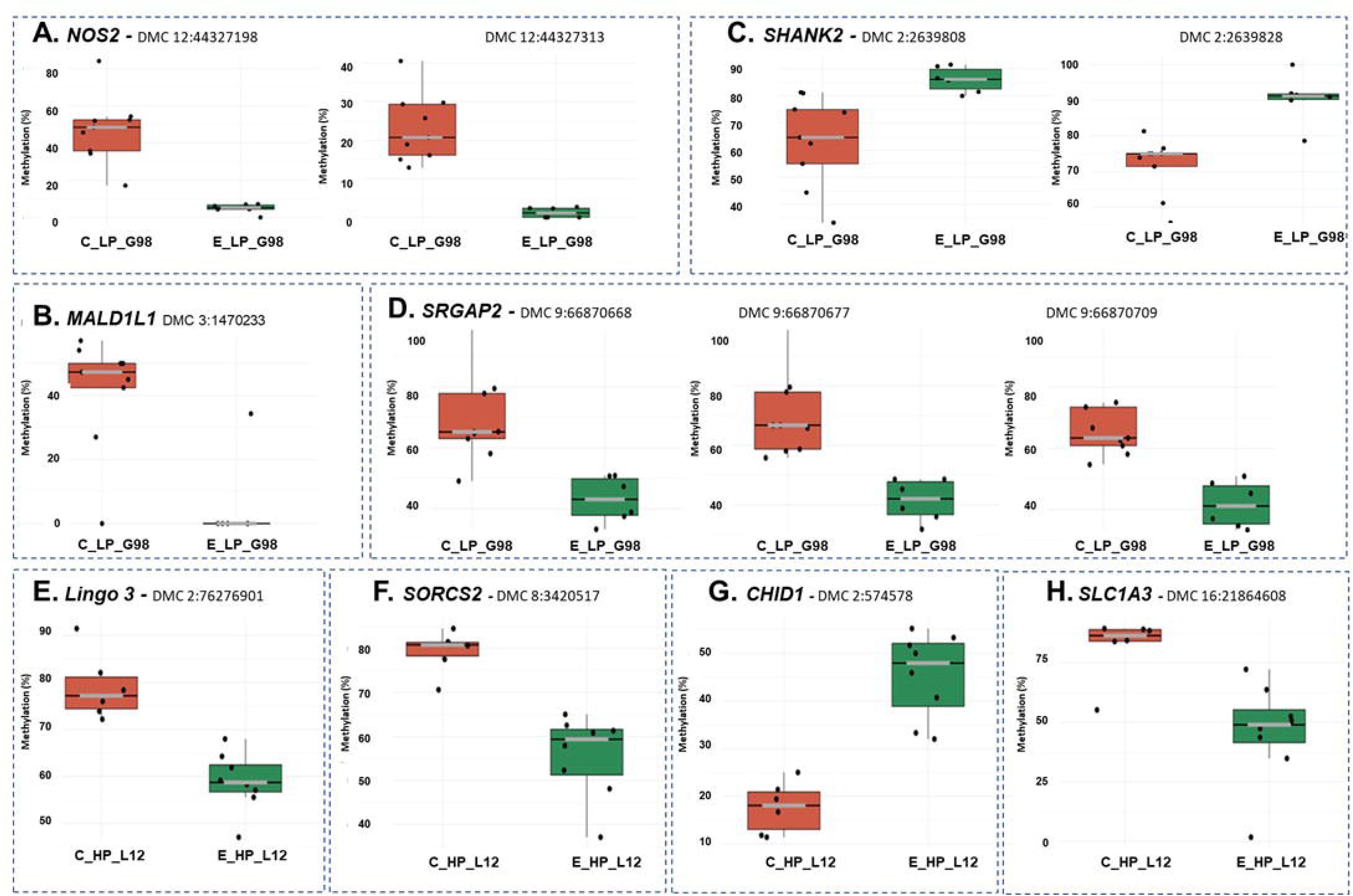
Methylation level of housing related DMCs targeting different genes. (**A**) *NOS2* – 2 DMCs; (**B**) *MALD1L1* – 1 DMC; (**C**) *SHANK2* – 2 DMCs; (**D**) *SRGAP2* – 3 DMCs; (**E**) *LINGO3* – 1 DMC; (**F**) *SORCS2* – 1 DMC; (**G**) *CHID1* - 1 DMC; (**H**) *SLC1A3* – 1 DMC. They were identified either in LP group at G98 (A, B, C and D) or in HP at L12 (E, F, G and H).

**Table 5:** Genomic distribution of differentially methylated cytosines (DMCs) in enriched (E) sows relative to conventional (C) sows in low-parity (LP) or high-parity (HP) groups on gestation day 98 (G98) and lactation day 12 (L12). ^1^Numbers and calculated percentages are indicated; percentages were based on the total numbers of DMCs at each time point. ^2^As a reference, the genomic distribution of 1,433,115 CpG_10-500_ (CpG_10-500_ background) was determined. The box colour indicates whether a preferentially genomic distribution of DMCs was observed compared to that of CpG_10-500_ background (green = increase and red = decrease).

|  | LP sows |  | HP sows |  | CpG<sup>10-500</sup><br>background<sup>2</sup> |
| --- | --- | --- | --- | --- | --- |
|  | DMCs G98<sup>1</sup> | DMCs L12<sup>1</sup> | DMCs G98<sup>1</sup> | DMCs L12<sup>1</sup> |  |
| CpG density in gene bodies versus intergenic regions |  |  |  |  |  |
| Intergenic | 23<br>(38.3%) | 14<br>(40%) | 17<br>(40.4%) | 23<br>(28.3%) | 284,740<br>(19.8%) |
| Exon | 5<br>(8.3%) | 4<br>(11.4%) | 2<br>(4.7%) | 6<br>(7.4%) | 228,050<br>(15.9%) |
| Intron | 25<br>(41.6%) | 13<br>(37.1%) | 19<br>(45.2%) | 36<br>(44.4%) | 479,396<br>(33.45%) |
| Promoter | 2<br>(3.3%) | 0<br>(0%) | 0<br>(0%) | 3<br>(3.7%) | 134,874<br>(9.4%) |
| TSS | 1<br>(1.6%) | 1<br>(2.8%) | 0<br>(0%) | 0<br>(0%) | 83,225<br>(5.8%) |
| TTS | 0<br>(0%) | 0<br>(0%) | 0<br>(0%) | 0<br>(0%) | 5,562<br>(0.3%) |
| 3'UTR | 2<br>(3.3%) | 1<br>(2.8%) | 1<br>(2.3%) | 7<br>(8.6%) | 51,146<br>(3.5%) |
| 5'UTR | 2<br>(3.3%) | 2<br>(5.7%) | 3<br>(7.1%) | 6<br>(7.4%) | 166,120<br>(11.5%) |
| CpG density associated with CpG islands |  |  |  |  |  |
| CpG desert | 38<br>(63.3%) | 29<br>(82.8%) | 29<br>(69%) | 56<br>(69.1%) | 494,291<br>(34.4%) |
| CpG Island | 8<br>(13.3%) | 0<br>(0%) | 4<br>(9.5%) | 13<br>(16%) | 701,098<br>(48.9%) |
| CGi shelves | 4<br>(6.6%) | 2<br>(5.7%) | 4<br>(9.5%) | 3<br>(3.7%) | 86,408<br>(6%) |
| CGi shore | 10<br>(16.6%) | 4<br>(11.4%) | 5<br>(11.9%) | 9<br>(11.1%) | 151,316<br>(10.5%) |
| Overlapping with repeat sequences |  |  |  |  |  |
| Without overlapping | 41<br>(68.3%) | 20<br>(57.2%) | 29<br>(69.1%) | 49<br>(60.4%) | 1,123,127<br>(78.36%) |
| With overlapping | 19<br>(31.7%) | 15<br>(42.8%) | 13<br>(30.9%) | 39<br>(39.6%) | 309,986<br>(21.64%) |

## Discussion

The aim of this study was to investigate how environmental enrichment throughout the reproductive life of pregnant sows affects DNA methylation profiles in immune cells. To do so, we compared PBMCs from sows of different parities and involved in either the gestation or the lactation phase of their reproductive cycle. In our previous research on the same animals, we demonstrated the positive effects of environmental enrichment on cortisol secretion, explorative and social behaviours of the sows, indicating an improved welfare^27^. In the same study, some minor housing effects on gene expression in their PBMCs were reported without significant alteration of blood formula^28^. This additional study has introduced analytical approaches designed to differentiate between true environmental effects and those linked to the reproductive life of animals, while minimizing potential biases arising from individual genetic background.

### Putative genetic influence on individual DNA methylation profiles

Some of our results suggested a genetic influence on inter-individual variations in DNA methylation profiles. In humans, growing evidence has shown that genetic variations play a role in the establishment of DNA methylation marks both in *cis* (same chromosome) and in *trans* (across chromosome^39,40^). In our study, genetic diversity could be expected because the experiment was performed at a commercial farm using sows resulting from Landrace x Large White crosses and purchased from various breeder, without animal genotyping. In order to limit genetic influence, data filtering was applied using two lists of known SNPs in the porcine species^33,35^. This filtering only concerns the overlapping SNP/CpG. Then, we considered for each DMC (and DMR), the distribution of methylation levels between individuals. However, we identified some DMCs with tri-modal distribution of methylation level, which may be imputable from individual genotype. A marked example of a putative genetic influence on our results was DMR with a gain of methylation identified in the HP group at the 5’UTR position of the *MYLK* gene, associated with a type I transposon and CGi shelf*. MYLK* encodes non-muscle myosin light chain kinase, which facilitates the interaction between myosin and actin filaments to produce contractile activity, particularly in neutrophils^41^. In our study, we observed a tri-modal distribution of methylation levels of *MYLK* DMR for different animals classified either in LP or HP groups. Most LP individuals exhibited intermediate DNA methylation levels, whereas HP individuals predominantly displayed high methylation levels, suggesting an apparent genetic effect on *MYLK* DMR methylation. In humans, previous research had found a strong association between the methylation levels of specific CpGs and the presence of SNPs in this gene^42^. In addition, Sun et al.^43^ demonstrated that the activity of *MYLK* promoters is influenced by factors such as LPS, TNFα, and hypoxia-inducible transcription factors, and is subject to strong epigenetic regulation, further modulated by proximal SNPs^43^. Our results suggest that genetic-epigenetic interaction at the *MYLK* promoter level, observed in humans, might be precisely studied in pig.

### Influence of parity class on individual methylome

The hierarchical clustering of the filtered background suggested a significant effect of parity class on individual DNA methylation profiles. Additionally, a significant loss of global methylation in HP sows was observed. According to this data, we identified a large number of DMCs (1,358 at G98 and 680 at L12) between high- and low-parity sows with and, consistently, more DMCs with a lower methylation level in HP sows. A higher proportion of DMCs affected by parity class were located within intronic regions. This does not typically obstruct transcription but can enhance gene expression by affecting regulatory elements such as enhancers and can also affect mRNA processing/alternative splicing^44,45^. Interestingly, contrary to the genomic distribution of all background CpG, most parity-associated DMCs were located in CGi deserts, regions typically associated with chromatin status and important for distal regulation of gene^46^. Most DMCs displayed a continue distribution of methylation levels, suggesting that they could be considered as epigenetic biomarkers of parity.

### Influence of physiological stage on parity-epigenetic effect

The reduced number of parity-associated DMCs observed at L12 in this study was consistent with our previous transcriptomic findings, showing a decrease of differentially expressed genes at this physiological stage when compared to the end of gestation^27^. Gestation and lactation are two periods highly hormonally different. Gestation is well known to be associated with maternal physiological adaptations to support foetal development. In particular, maternal leukocytes play a key role in managing inflammation and maintaining immune tolerance towards the foetus. The transition from gestation to lactation involves significant hormonal changes. These hormonal changes may support major evolution in immune competence and response^47,48^. We previously reported, using the same blood samples, a significant difference in monocyte and neutrophil numbers between G98 and L12 as well as in lymphocyte numbers in association with parity, without interactions between the two conditions (physiological stage x parity^27,28^). Epigenetic data (herein investigated) such as transcriptomic data^27^ were produced using the same PBMC preparations, rich in lymphocytes and monocytes. Taken together, these findings suggested that the substantial metabolic and endocrine reprogramming, as well as weak immune cell proportion variations that occurs during the gestation-lactation transition, may contribute to explain a part of DNA methylation variations.

### Identification of epigenetic markers associated with parity class

We identified numerous DMRs when comparing high- vs. low-parity sows, suggesting a consistent epigenetic effect on the genes concerned. Many DMRs co-localized with regulatory regions suggesting their involvement in gene expression regulation. After excluding DMRs with a tri-modal DNA methylation distribution, four parity DMR-associated genes were linked to the epigenetic machinery: *DNMT3A* (encoding DNA methyltransferase 3A), *KDM8* (encoding a histone demethylase that preferentially recognizes and cleaves monomethylated and demethylated arginine residues on histones H2, H3, and H4) at G98, *SIRT2* (encoding an NAD+ (nicotinamide adenine dinucleotide-dependent deacetylase) at L12, and *HDAC4* (encoding a class II histone deacetylase) common to both physiological stages. *DNMT3A-* DMR, located in an intron, exhibited a gain of methylation in the HP group and co-localized with an enhancer. DNMT3A, a crucial enzyme responsible for establishing de novo DNA methylation, plays a pivotal role in maintaining the delicate balance between hematopoietic stem cell differentiation and self-renewal^49^. In humans, DNMT3A function loss mediates changes in the epigenetic landscape and can promote aberrant lymphocyte differentiation and function^49^. In our study, it could be hypothesized that the methylation status of *DNMT3A*-DMR could induce a reduction of *DNMT3A* transcription, consistent with the global loss of methylation observed in the HP group. *KDM8*-DMR, located in exonic region, exhibited a gain of methylation in the HP group. KDM8 functions as a transcriptional activator by inhibiting HDAC (histone deacetylase) recruitment via the demethylation of H3K36me2. Additionally, *HDAC4*-DMR, located in an intronic region, exhibited a loss of methylation in the HP group and co-localized with an enhancer, potentially increasing *HDAC4* gene transcription. Together, the opposite methylation patterns of *KDM8*-DMR and *HDAC4*-DMR in HP suggested a convergence of effects towards an increase in the expression of *HDAC4* gene and potentially its activity. Thus, the increase in parity rank could result in deacetylation in favour of chromatin closure. In our sows, age and parity rank were strongly correlated. Thus, in HP sows, global methylation loss, which is a common characteristic of ageing, could be associated to the increase in chromatin instability, while transcriptional inhibition involving HDAC4 activity could be more localized at specific genes.

Several parity-DMRs were associated with genes involved in the immune response, including *CD2, CAMK4, CMIP, SECTM1, SEC14L1, SGK1, SKI, UMODL1, PACS1, CD5* and *TNFRSF1B.* Based on the genomic features, functional annotation/regulatory landscapes and DNA methylation status of the DMR, it could be hypothesized that their expression was affected by parity class. Positive transcriptional regulation could affect *CAMK4, CD2, CMIP, SECTM1, UMODL1*, and *TNFRSF1B* in HP sows. *CAMK4* belongs to the serine/threonine protein kinase family, has a limited tissue distribution including NK and T-cells. *CD2* encodes a surface co-stimulatory receptor present on T-cells. Co-stimulation is needed in addition to antigen-specific signalling through the T cell receptor CD3 complex to achieve full T-cell activation. CD2 expression is increased in memory T-cells, whose numbers increase with ageing^50^. *CMIP* is a negative regulator of CD28, another T-cell co-stimulatory molecule, whose expression and signalling decreases with ageing in T-cells^51^. It also leads to activation of the c-Maf Th2-specific factor^52^. *SECTM1* is primarily expressed in neutrophils and monocytes in peripheral blood. It acts as a ligand for the CD7 receptor, which is also a T-cell co-stimulatory receptor, and synergizes with suboptimal anti-CD28 to increase T cell functions^53^. *TNFRSF1B*, also known as Tumor necrosis factor (TNF) receptor 2, is expressed, among blood circulating immune cells, only by T-cells. TNFR2 is necessary for antigen-induced differentiation, for the survival of T cells, and is also involved in the regulation of the inflammatory response^54^.

In contrast, negative transcriptional regulation could affect the following genes in HP sows: *SEC14L1, SGK1, SKI, and PACS1. SEC14L1* encodes a protein that negatively regulates *RIG-I*^55^, an intra-cytosolic receptor expressed ubiquitously that recognizes RNA viruses and triggers a signalling cascade resulting in the production of type I interferons and pro-inflammatory cytokines^56^. *SGK1* is a serine/threonine protein kinase that activates certain potassium, sodium and chloride channels. It is upregulated by cell stress, inflammation as well as a variety of hormones including glucocorticoids^57^. In T-cells, SGK1 acts for differentiation towards the pro-inflammatory Th17 lineage and inhibits development towards the T regulatory phenotype^58^. SKI is a nuclear proto-oncogene that represses TGF-beta signalling, a ubiquitous cytokine acting as a negative growth regulator and playing a major role in the negative regulation of immune responses^59^. *PACS1*, a highly conserved cytosolic protein that facilitates the trafficking of cargo between membrane-bound compartments. It is known to be expressed in human peripheral blood leukocytes and is important for maintaining peripheral T and B-cell populations and favours lymphocyte quiescence and survival^60^.

Comparisons with the transcriptomes of the same PBMCs used in the present study^27^ revealed 28 genes affected at both the epigenetic and transcriptome levels at G98 (*AOC1, ANXA2, BCL3, CHST1, CUX2, DIP2C, DNMT3A, DSP, EPB41L5, EXOC7, GRAMD4, IL9R, JUP, LDLRAD4, MPV17, PRDM11, PRKAR1B, RCHY1, RUNX2, SLC22A15, SLC6A17, SORCS2, SOX13, SYTL3, TATDN3, TNS1, TNFRSF4,* and *URB1*). Both transcriptome and methylome data agreed regarding the influence of parity class on biological processes (BP) such as cell-cell adhesion functions (notably with the *ANXA2* gene affected in both datasets), the ERK1/ERK2 cascade, and cell surface signalling pathways (particularly those involving G protein-coupled receptors in both datasets). The effects of parity on T-cell differentiation (with the *RUNX2* gene affected in both datasets), positive regulation of INF-γ (notably with the *BCL3* gene in both datasets) and the positive or negative regulation of transcription also appeared in both datasets, either during gestation or lactation. Previous works in the pig had demonstrated immune differences between primiparous and multiparous animals^61,62^, but the effects of successive reproductive cycles in multiparous females were not investigated. Taken together, our results indicate that among multiparous sows, parity class remains an important factor that should be considered when exploring immune-related characteristics.

The parity rank was highly correlated with the sows’ age. In humans, age is a well-known factor that induces a loss of DNA methylation across various cell types and tissues^63^. Epigenetic aging can be assessed using various epigenetic clues. Horvath proposed a method to measure the cumulative effect of an epigenetic maintenance system throughout an individual’s life (from various tissues, including PBMCs)^64^. A similar study was conducted in pigs^25^. In our study, we identified 13 genes that were common to both Horvath’s list and our list containing the 1,786 DMC-associated genes affected by parity at both time-points (*DST, FES, GPR68, GRIN2C, LOXL2, MN1, NDUFA13, RAPGEF1, RXRA, SCD5, SLC28A2, ST3GAL4* and *UROS*). However, we believe that the differences observed between parity classes could not solely be linked to the aging process. The sows in this study could not be considered as old because their ages ranged from 16 to 36 months, while the natural average lifespan of a pig is approximately 15 years. The possibility of a parity effect (i.e. of accumulated reproductive cycles and not of aging) on DNA methylation is supported by data in humans. Some studies in women have suggested that parity either slows down blood epigenetic aging^65–68^ or accelerates cell senescence, epigenetic ageing and a shortening of telomeres^69–71^.

### Epigenetic marks of environmental enrichment

Despite a clear effect of environmental enrichment on behaviour and salivary cortisol levels in the sows used in this study^27,28^, suggesting their improved welfare, the effects of housing environment on DNA methylation were modest. This weak epigenetic effect was in line with our previous transcriptomic observations in PBMC, with few genes affected by the housing environment^27^. Environmental enrichment generated DMCs in genomic regions similar to those seen for parity, i.e. mainly in intronic regions and CpG desert, which might influence chromatin structure and accessibility. Our results agreed with those of a previous study which showed that environmental enrichment increased chromatin accessibility and transcription factor binding in the brains of young mice^72^.

In both parity groups, environmental enrichment modulated the methylation of a few DMC-associated genes involved in the emotion functions or disorders resulting from exposure to severe stressors in humans, making these genes relevant candidates for assessing affective states in pigs. In the low-parity group, sows housed in the enriched environment exhibited a loss of methylation in two DMCs in the intronic regions of the *SHANK2* gene located inside a CpG island, and in one DMC in the intronic region of the *MAD1L1* gene. *SHANK2* encodes a synaptic protein. In human, its aberrant expression has been implicated in phenotypes such as intellectual disability and depression^73^. Its role in immune cells is unknown but gain of methylation of the *SHANK2* promoter region has been observed in the blood of humans exposed to severe life stressors^74^. In pig, five different *SHANK2* transcripts were previously described with variations in coding exons leading to different protein isoforms. Further molecular investigations are needed to be explore the functional epigenetic change induced by environmental enrichment. In humans, specific DNA methylation profiles of *MAD1L1* were shown to be linked to reward system functioning, anxiety regulation and associated psychiatric diseases^75,76^. In human immune cells, methylation variations of this gene throughout life were also associated with depression^77^, and a higher gene expression was reported in self-reported highly stressed individuals^78^.

In the high-parity group, environmental enrichment was associated with a loss of methylation compared to conventional control sows in an exonic CpG in *LINGO3* gene, and an intronic CpG in *SORCS2* gene. *LINGO3* encodes a protein involved in the myelinisation process and has been linked to depression^79^. *SORCS2* codes for a receptor of brain-derived neurotrophic factor (BDNF), which is involved in synaptic plasticity^80^ and the regulation of mood disorders^81^. In mice, environmental enrichment increased BDNF levels in the brain through epigenetic processes^82^. In pigs, environmental enrichment increased BDNF plasma levels, which have been proposed as an indicator of animal welfare^83^. Three other genes for neurological functioning, regardless of parity class and time point, were identified (*SRGAP2, GRIK4, SLC1A3*). *SRGAP2*, coding SLIT-ROBO Rho GTPase–activating protein 2, is involved in controlling human brain plasticity^84^, but also in cell polarization, regulating neutrophils firm adhesion to endothelial cells^85^, and consequently playing a critical role in inflammatory responses. The loss of methylation observed in an intronic CpG of *SRGAP2* may be associated with a downregulation of its expression. *GRIK4* encodes a protein that belongs to the glutamate-gated ionic channel family. Glutamate functions as the major excitatory neurotransmitter in the central nervous and glutamate dysfunction is associated mood disorders^86^. *SLC1A3,* encodes a member of a high affinity glutamate transporter family which plays role in the termination of excitatory neurotransmission. It is also associated with major depressive disorders in humans^87^. Thus, the loss of methylation of a CpG detected herein in the 5’UTR region of *GRIK4* gene, as well as in the CpG detected upstream *SLC1A3* gene, may be associated with up-regulation of their expression, putatively affecting the levels of the excitatory signal of glutamate in the sows. Collectively, our results highlight some interesting biomarkers of environment effect, occurring in peripheral blood cells, but with putative impact on brain activity and mood regulation. The positive effects of environmental enrichment on brain structure and function have been demonstrated extensively in the literature on rodents^88^. Some studies have also explored the relationship between DNA methylation in brain regions and environmental enrichment. For example, Zocher et al. compared aged and young mice housed in enriched conditions and found that environmental enrichment preserved a methylation pattern in aged mice similar to that seen in younger individuals^89^.

## Conclusion

This study suggested a marked influence of animal genetic background on individual DNA methylation profile, emphasizing the need for a cautious interpretation of the results. Despite this putative genetic influence, a significant effect of parity was observed, even among multiparous sows. This highlights the importance of reproductive life history as a key factor in modulating epigenetic mechanisms related to immune cell function in female animals. Pigs could therefore be considered an effective model for investigating the impact of successive gestation and lactation cycles on maternal DNA methylation profiles and for exploring potential shifts in the epigenetic clock. Furthermore, our findings indicate that positive environmental enrichment exerts a modest effect on DNA methylation profile. This suggests that while environmental enrichment may have beneficial effects on behavioural and physiological aspects, these benefits may be translated into limited molecular changes. Nevertheless, this study provides an original set of biomarkers that need to be further validated and could be routinely followed in further studies with diversified environments in pig farms.

## Supporting information

Suppl Table 1_Sheets 1-6:

Suppl-Table 2 Global methylation analysis using SNP_filtered 1 433 115 CpGs

Suppl_Table3 Functional Annotation of parity related DMRs

Suppl_Table 4 Functional Annotation of housing related DMCs

## Ethics of Experimentation

The experiment was carried out at the Chambre Régionale d’Agriculture de Bretagne (CRAB) Experimental Farm in Crécom (Saint - Nicolas du Pélem, France), and complied with EU Directive 2010/63/EU on the protection of animals used for scientific purposes. It was approved by the French Regional Ethics Committee on Animal Experiments no 007 and by the French Ministry of Higher Education and Research (APAFIS#27038-2020090315423669 v2). Moreover, this study adheres to ARRIVE guidelines.

## Data availability

The data supporting the findings of this study will be available from the corresponding author upon reasonable request until December 31, 2025. Thereafter, the raw and processed RRBS datasets will be publicly accessible at the European Nucleotide Archive (ENA) under project accession PRJEB90105 (secondary accession ERP173119).

## Supplementary materials availability

All supplementary figures, tables and data sets are upload to a public repository and available using the link “, https://doi.org/10.57745/0GWNEV, Recherche Data Gouv. and referenced under “MERLOT, Elodie; JAMMES, Hélène, 2025, “BEPPI project – data, tables and figures on sow peripheral white blood cell epigenome”.

## Authors contribution

M. M. Lopes: Data curation, formal analysis, Investigation, validation, visualization, writing original draft. G. Monteiro Moreira: Resources, visualization, writing review editing. A. Chaulot. Talmon.: investigation, resources. A. Frambourg.: Resources, software, writing review editing. V. Costes: Resources, software, writing review editing. J. Demars: Resources, writing review editing. E. Merlot: conceptualization, funding acquisition, investigation, project administration, supervision, writing review and editing. H. Jammes: conceptualization, supervision, visualization, writing review and editing

## Acknowledgments

The authors are grateful to C Gerard and N Villain, engineers from Brittany’s Regional Agricultural Chamber (CRAB), and C Connan and P Lirzin, staff at the CRAB Experimental Farm, for their help in monitoring the experiment, sampling and data collection. We also thank C Trombani, veterinarian (Breizh Pig, France), for his assistance during blood sampling procedures. We would also like to acknowledge A Vincent, F Thomas, L Le Normand, and P Touanel for their technical assistance (PEGASE, INRAE, Saint-Gilles, France). Our thanks also go to the sows who took part in this study.

## Supplemental material

**Suppl. Figure 1.** Hierarchical non-supervised clustering using a matrix obtained before SNP filtering and including the methylation level of 3,418,048 CpGs_10-500_ for all libraries (n = 58). The library names are coloured according to (**A**) housing conditions and to (**B**) maternal genealogy (Abbreviations: C, Conventional housing; E, enriched housing; DM, different mother; MHS, maternal half-sisters, as described in Table 1). The bar underlines the libraries produced from the sows included in the two replicates (n=8).

**Suppl. Figure 2.** Hierarchical non-supervised hierarchical clustering using a matrix obtained after SNP filtering and including the methylation level of 1,433,115 CpGs_10-500_ for all libraries (n = 58). The library names were colored according to (A) assay design and to (B) paternal genealogy. Abbreviations: R, replicate; DF, different father; P-HS, paternal half-sisters - described in Table 1). The bar underlines the libraries produced from the sows included in the two replicates (n=8).

**Suppl. Figure 3.** (**A**) Genome-wide distribution of 1,433,115 CpGs_10-500_, analysed by RRBS, SNP filtered and common at all libraries. (**B**) Genome-wide distribution of parity related-DMCs at L12. Pig 18 autosomes and X chromosomes are represented by horizontal bars according to their size (https://www.ncbi.nlm.nih.gov/datasets/genome/GCF_000003025.6/). DMCs identified between low and high parity on L12 are represented by coloured thin lines: blue lines for less methylated DMCs and red lines for more methylated DMCs in the HP group.

**Suppl. Figure 4.** For each DMR, the chromosomic position and sequence of DMR and gene organization were represented. The individual methylation rates for each CpG (named by the start genomic position) are shown as a bar chart for each individual from LP (in pink) and HP (in red) groups at G98 and L12. The correlations between methylation levels at G98 and L12 were presented in the LP group (in pink) and in the HP group (in red). **(A)** SGK1 – DMR1 (located on chromosome 1, 3 DMCs), **(B)** SGK1 – DMR2 (located on chromosome 1, 5 DMCs) and **(C)** HDACA – DMR (located on chromosome 15, 3 DMCs).

**Supplementary Table 1_Sheet 1**: Overview of sequencing parameters for all samples

**Supplementary Table 1_Sheet 2**: Differentially methylated cytosines (DMCs) affected by parity on gestation day 98 (G98)

**Supplementary Table 1_Sheet 3**: Differentially methylated cytosines (DMCs) affected by parity on lactation day 12 (L12)

**Supplementary Table 1_Sheet 4**: Differentially methylated regions (DMRs) affected by parity on gestation day 98 (G98) and lactation day 12 (L12)

**Supplementary Table 1_Sheet 5**: Differentially methylated cytosines (DMCs) affected by environmental enrichment in low-parity sows on gestation day 98 and lactation day 12

**Supplementary Table 1_Sheet 6**: Differentially methylated cytosines (DMCs) affected by housing conditions in high-parity sows on gestation day 98 and lactation day 12

**Supplementary Table 2**: Global methylation analysis using SNP_filtered 1 433 115 CpGs

**Supplementary Table 3**: Functional Annotation of parity related DMRs

**Supplementary Table 4**: Functional Annotation of housing related DMCs

**Supplementary data 1**: Methylation distribution_common_parity_DMCs

**Supplementary data 2**: Methylation distribution_parity_gestation_DMRsall

**Supplementary data 3**: Methylation distribution_parity_lactation_DMRsall

**Supplementary data 4**: Methylation distribution_housing_LP_G98_DMCs

**Supplementary data 5**: Methylation distribution_housing_LP_L12_DMCs

**Supplementary data 6**: Methylation distribution_housing_HP_G98_DMCs

**Supplementary data 7**: Methylation distribution_housing_HP_L12_DMCs

