## Supplementary figures and images for "The effect of environmental enrichment on immune cell DNA methylation profiles depends on the parity of sows"

### Suppl_Figure4B

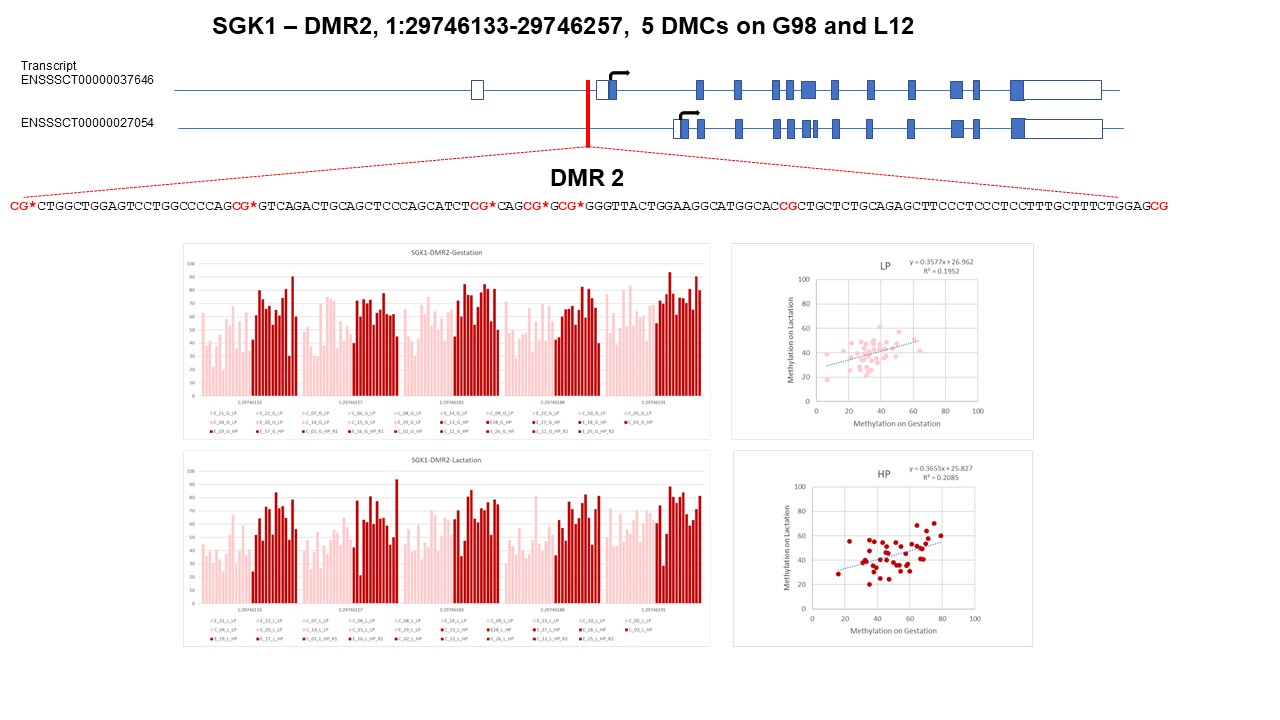

### Suppl_Figure 1

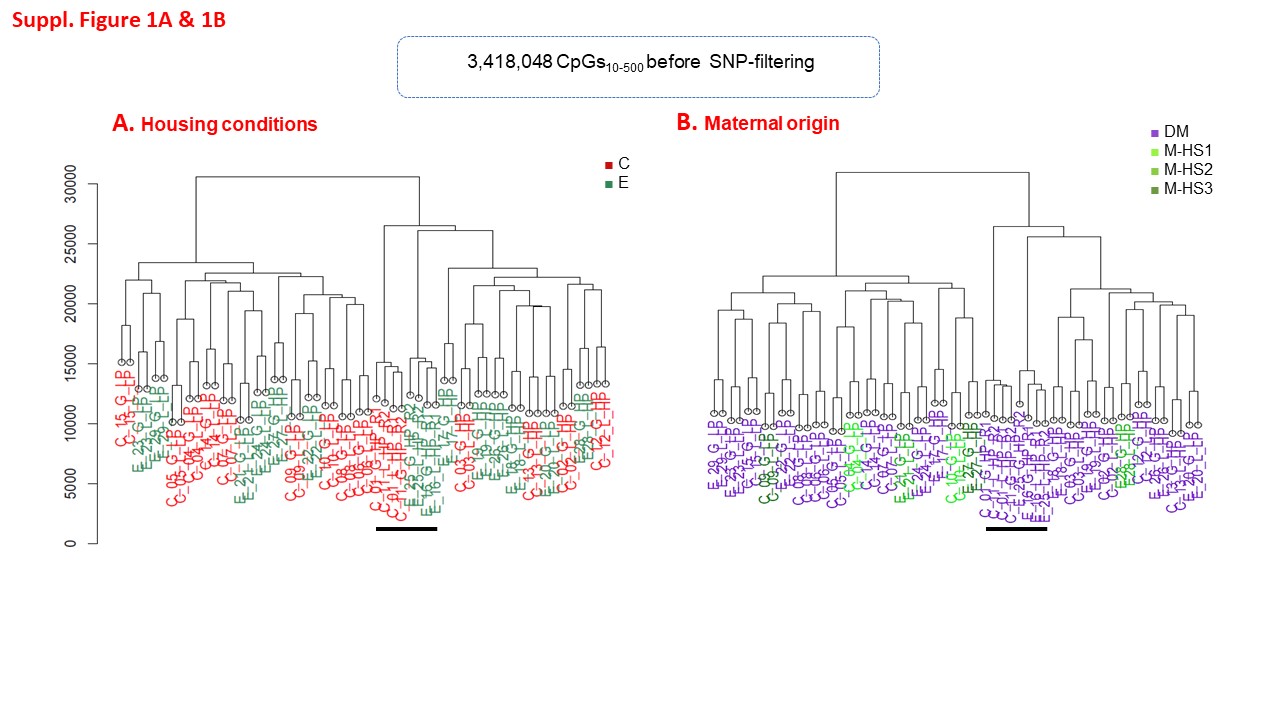

### Suppl_Figure 2

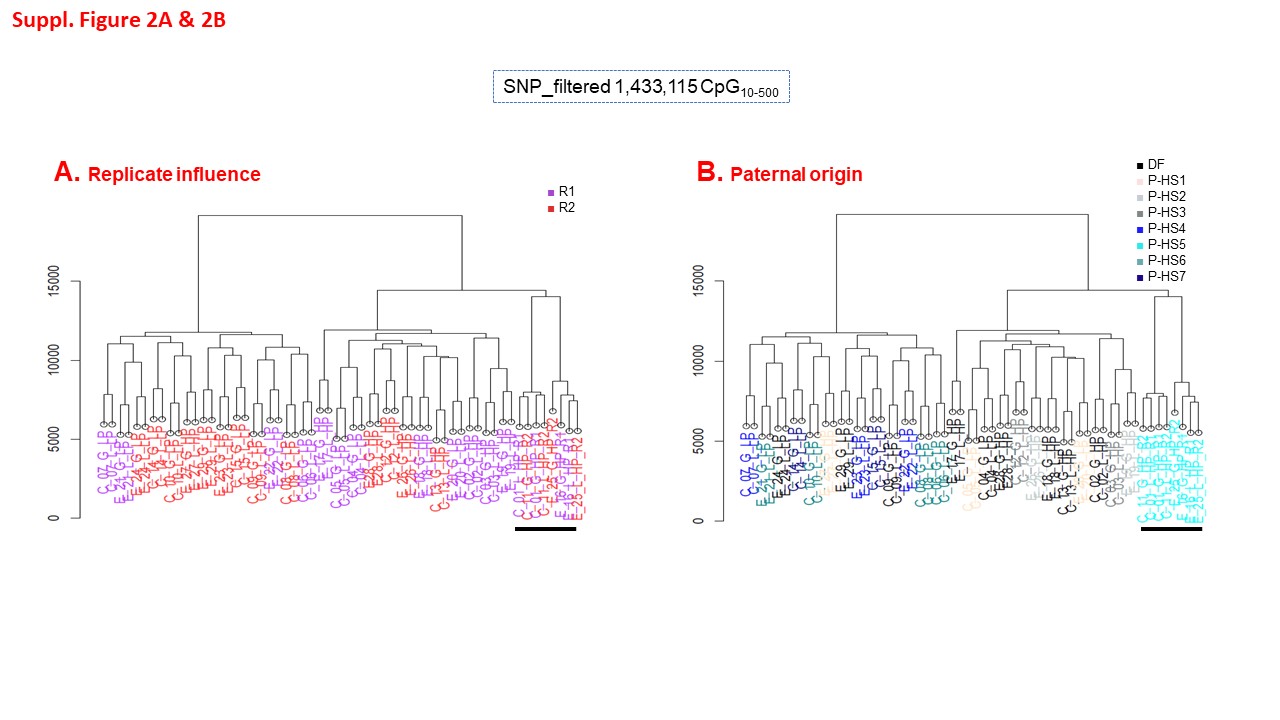

### Suppl_Figure 3A

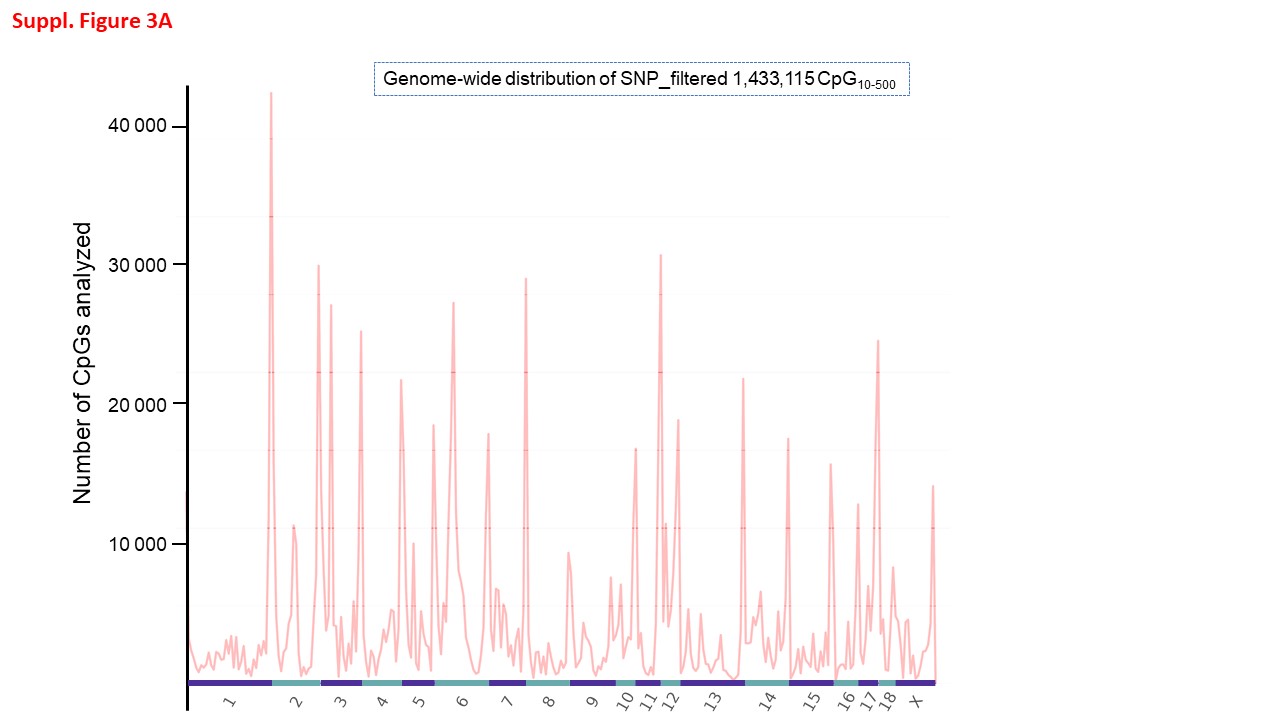

### Suppl_Figure 3B

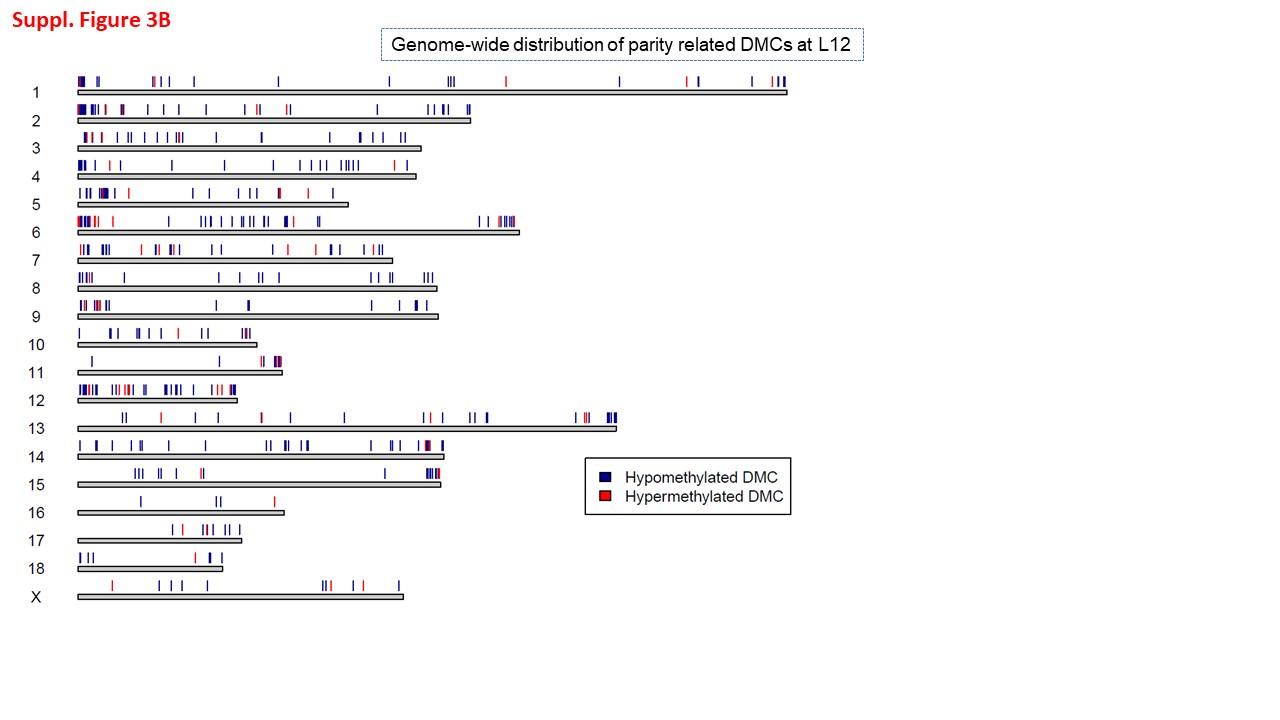

### Suppl_Figure 4A

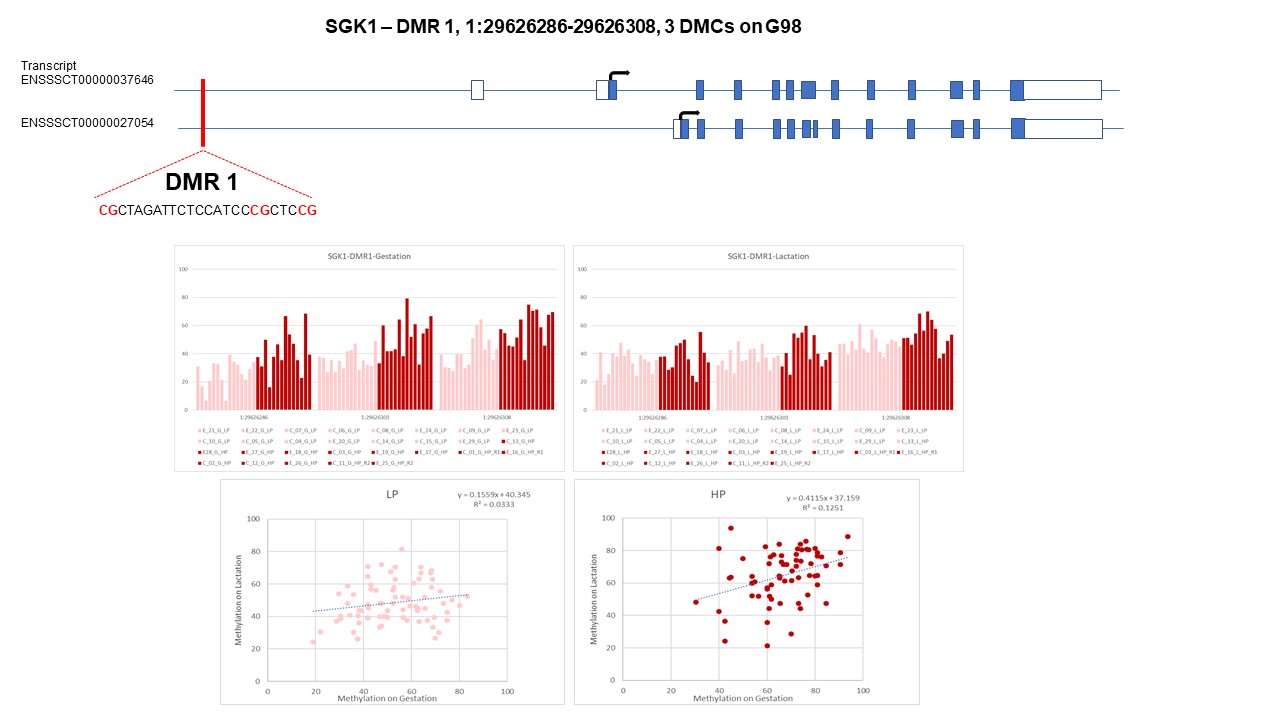

### Suppl_Figure 4C

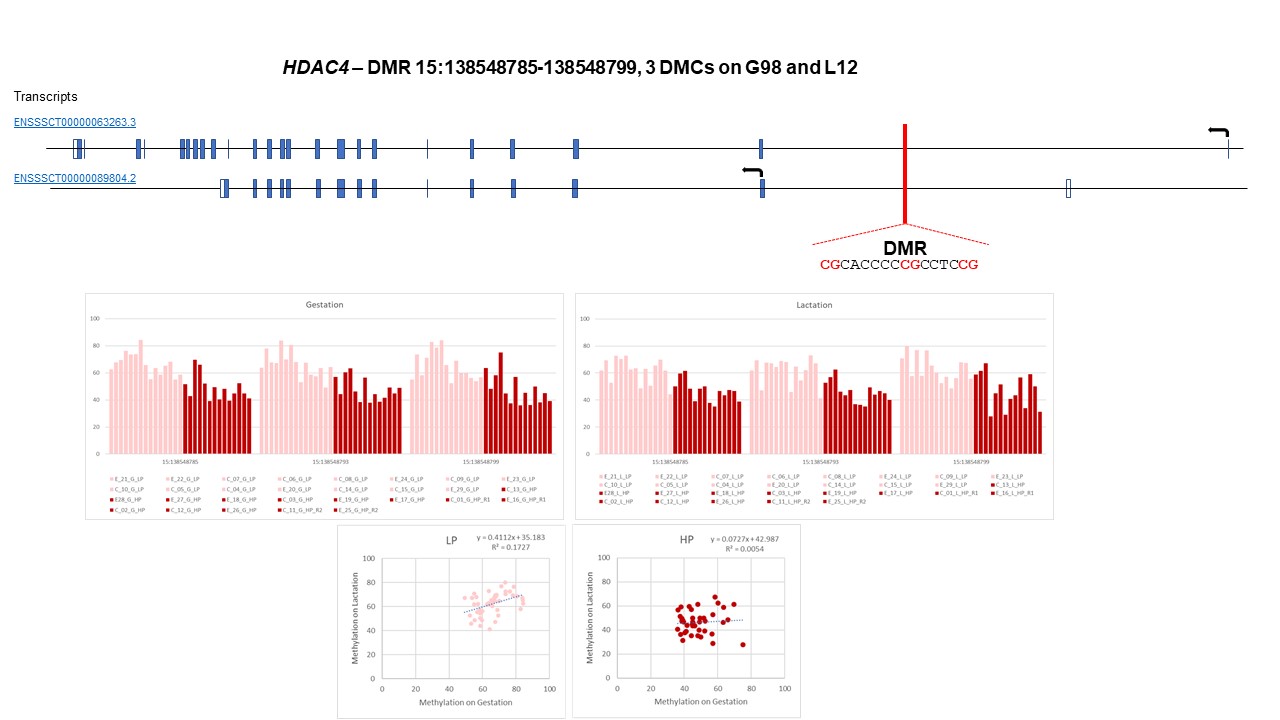
